# Supervised Deep Learning for Efficient Cryo-EM Image Alignment in Drug Discovery with cryoPARES

**DOI:** 10.1101/2025.03.04.641536

**Authors:** Ruben Sanchez-Garcia, Alex Berndt, Amir Apelbaum, Judith Reeks, Pamela A Williams, Carl Poelking, Charlotte M Deane, Michael Saur

**Author notes:** These authors jointly supervised this work (Charlotte M Deane, Michael Saur).

## Abstract

Cryo-Electron Microscopy (cryo-EM) is a pivotal tool for determining 3D structures of biological macromolecules. Current workflows are computationally demanding and require manual intervention, creating bottlenecks for high-throughput applications like structure-based drug discovery. In such contexts, where all protein samples can be assumed to be equivalent at resolutions relevant for image alignment, information about particle poses from previous refinements could be reused. Existing methods, however, ignore this prior knowledge, aligning each dataset from scratch. We present cryoPARES, a deep learning pose estimation method trained on pre-aligned datasets. Our method not only provides accurate angular predictions significantly faster than traditional approaches but also introduces automated particle pruning capabilities that eliminate manual intervention. Together with its single-pass operation, these features enable near real-time reconstructions that provide feedback during data acquisition. We demonstrate cryoPARES’s effectiveness through rapid structural determination of seven ligand-bound complexes across four distinct protein targets. We also release three fragment-bound cryo-EM datasets.

## 1. Introduction

Cryo-Electron Microscopy (cryo-EM) has emerged as a powerful tool in structural biology^1^ with applications in structure-based drug discovery (SBDD)^2–4^. At the heart of this technique lies Single-Particle Analysis (SPA), which reconstructs 3D structures of macromolecules from hundreds of thousands of 2D projection images, termed particles. Ideally, these particles adopt random orientations in the ice layer^5^, with their orientation initially unknown. The main challenge in SPA lies in estimating these orientations amidst high noise and multiple sources of error.

Typically, the orientation estimation is performed via iterative angular refinement^6–12^, which begins with a low-resolution reference volume against which each particle is compared in all possible orientations. The best-matching orientation(s) are then assigned to each particle, and a new volume is then reconstructed, serving as the reference for the next iteration. This process repeats until convergence. While powerful, this approach is computationally demanding due to the large number of iterations required and to the large number of comparisons performed in each iteration^13^.

For novel samples, the initial reference volume must be estimated ab initio^7,14–16^. However, when prior knowledge exists, it can be used to generate an initial volume. For instance, a previously refined map or a map simulated from an atomic model from a homologous protein can be used. While this strategy may introduce template bias^17,18^, using a higher resolution initial volume that is closer to the final solution can improve convergence of refinement algorithms and speed processing, although gains may be limited by the need for denser angular sampling at higher resolutions.

The use of already solved volumes as references has proven particularly successful in several situations. For example, in cases where it can be assumed that the overall protein structure remains largely unchanged upon ligand binding, a high-resolution structure of the apoprotein can serve as an excellent initial model for the determination of ligand-bound structures^19^. This is because pose estimation is mostly driven by low- to medium-resolution frequencies^20^ at which similar proteins are nearly identical. An extreme application of this idea is employed in the baited reconstruction by template-matching approach^19^, where a high-resolution reference obtained from an unbound atomic model was used to identify bound ligands via template-matching.

Despite these examples, the computational burden of processing multiple datasets of similar samples still limits wider adoption of cryo-EM in high-throughput studies. While high-resolution apo-structures can be leveraged for grid-based SO(3) searches, the dense angular sampling required for such searches typically imposes a significant computational bottleneck. Traditional hierarchical algorithms often perform a vast amount of redundant searching to avoid local minima, limiting the practical speed gains even when high-resolution prior information is available.

Moreover, current approaches require substantial manual intervention for parameter optimization and particle selection, creating additional bottlenecks in the pipeline. To address these challenges, several deep learning alternatives have been proposed^13,21–25^. Among these, “amortised pose inference” approaches, where a neural network is trained to predict particle poses directly from images, are particularly promising. While some are unsupervised ab initio approaches^26,27^, others can leverage previous structural knowledge through supervised learning^23–25,28^.

Supervised approaches train surrogate models of the alignment algorithm using pre-computed poses from previous refinements. While requiring pre-aligned particles may seem a limitation for ab initio refinement, this is less problematic in many applications, including ligand screening. In this context, the same protein target is repeatedly screened against different small molecule ligands, generating multiple datasets of previously aligned particles that can be used to train a model that is expected to generalize to similar particles bound to different ligands. However, none of these methods has yet demonstrated practical applicability in this scenario.

Building upon our previous work^28^, we present here cryoPARES (<u>cryo</u>-EM <u>P</u>ose <u>A</u>ssignment for <u>R</u>elated <u>E</u>xperiments via <u>S</u>upervision), a fully automatic pose estimation tool optimized for ligand screening. A single model trained on an apo structure can be used to efficiently process multiple ligand-bound datasets without manual intervention, demonstrating robust generalization across diverse samples. Furthermore, our tool performs automated particle pruning, eliminating the need for manual intervention in particle selection. It also shows promise in alleviating the preferred orientations problem often observed in cryo-EM samples. We validate these capabilities through the rapid structural determination of seven ligand-bound complexes across four distinct protein targets and release three new ligand-bound cryo-EM datasets to the community.

## 2. Results

### 2.1. CryoPARES is a tool to perform fast structural determination of related samples

CryoPARES addresses an increasingly common challenge in cryo-EM: processing multiple, closely related datasets. It is particularly valuable in SBDD, where work often proceeds only after at least one structure of the target protein has been resolved, and where many similar ligand-bound complexes are subsequently determined in succession. By reusing alignment information across these related experiments, cryoPARES minimizes redundant computation and accelerates structural determination.

To achieve this, cryoPARES uses a supervised deep-learning model trained on pre-aligned particles to predict the probability distribution of particle orientations. Once trained, the network rapidly determines poses for new, related datasets, yielding orientations accurate enough for high-resolution reconstruction. Contrary to conventional refinement methods, most of the computational cost occurs during the initial training (typically < 4 GPU days, see Supplementary Table 8), while subsequent analyses of similar specimens can be completed within minutes, greatly reducing the burden of large-scale structural studies (see Fig. 1). Although high-resolution reference volumes can be leveraged for traditional refinement algorithms (e.g., baited template matching), these approaches remain orders of magnitude less efficient than the amortized inference of cryoPARES due to the massive computational overhead of the orientation search over the dense orientation grids required for high-resolution accuracy (see Supplementary Note 5 and Supplementary Tables 6-8).

**Fig. 1.**
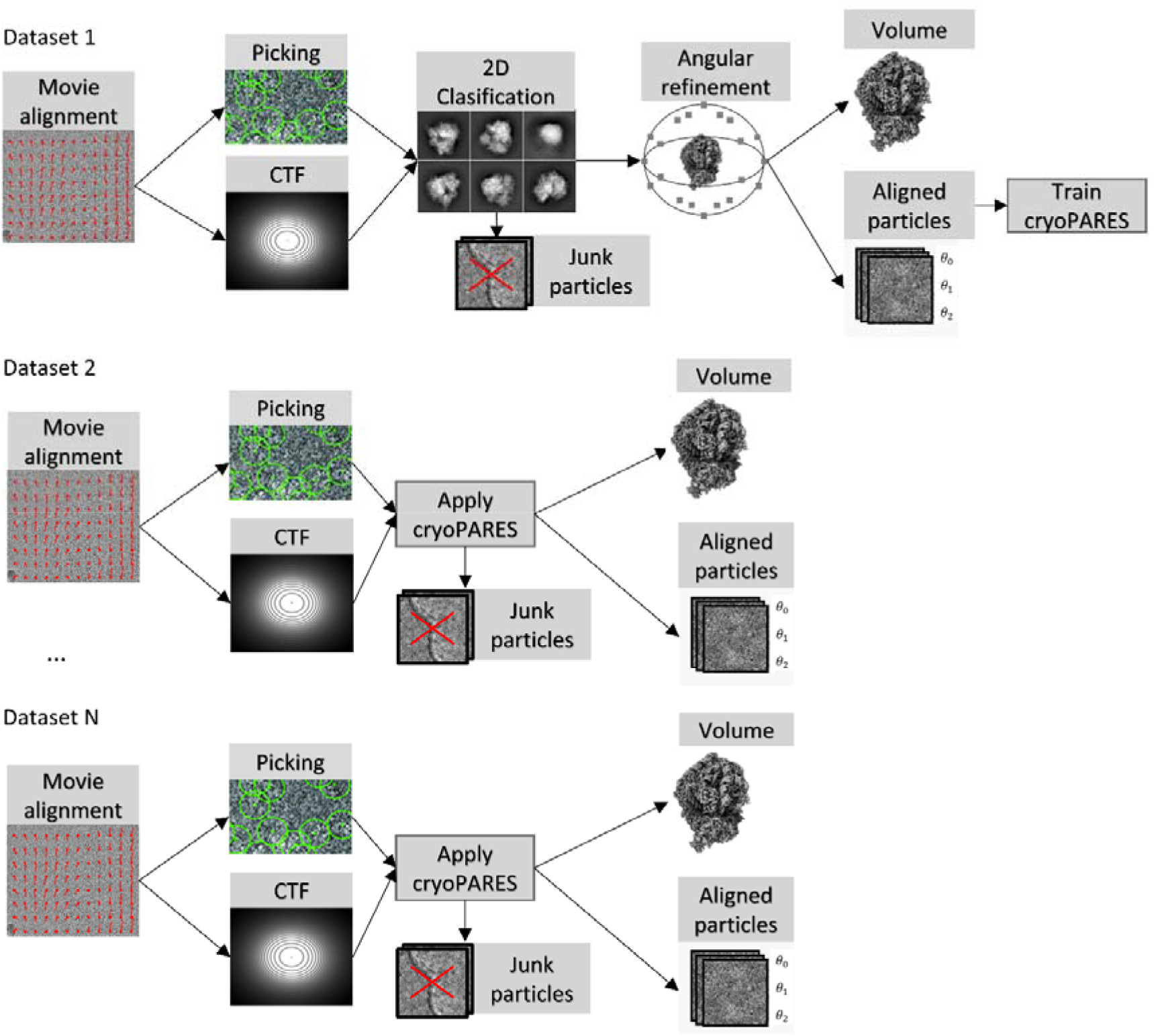
CryoPARES models are trained from a pre-aligned “good” particles dataset that is obtained via classical image processing pipelines. The model is trained in a supervised manner to predict the probability distributions of the pose of each particle. The junk particles in the training dataset can be used to calibrate the particle goodness score (see Methods Section 4.10). Once the models are trained, cryoPARES performs efficient structural determination of subsequent samples by replacing some of the most time-consuming steps by a quick one-shot estimation step. *θ*_*i*_ represents the alignment parameters of the particles.

Because cryoPARES predicts pose probabilities, it can automatically identify and exclude junk particles (see Results Section 2.5). This allows its direct use immediately after particle extraction, enabling a fully automated and reproducible reconstruction workflow while removing subjective steps such as manual 2D classification. Its one-shot inference, akin to baited reconstruction, determines orientations in a single pass and, together with its high processing speed (see Results Section 2.4), makes real-time image processing feasible, providing immediate feedback during data acquisition. Such automation and standardization are essential for high-throughput structural studies that demand speed, consistency, and reproducibility.

### 2.2. CryoPARES can resolve the structure of ligands bound to different protein targets

To illustrate the utility of our tool, we employed cryoPARES to determine the structure of several protein-ligand complexes for four different protein targets. In all cases, the cryoPARES models were trained using the pre-aligned particles of the apoprotein previously solved and a 6 Å low-pass-filtered apo map was used as the reference for local refinement.

We first tested our method on the previously published datasets of *E. coli* β-galactosidase (BGAL) bound to Compounds B1 and B2^3^. Fig. 2 shows the reconstructed densities of the ligand-bound datasets (c,d) compared with the apo reference used for training (a,b). The map features of the ligands using the orientations estimated with cryoPARES agree with the ground-truth atomic models reported previously. As a negative control, it can be observed that the densities of the ligands are not present in the apoprotein maps (Fig. 2 a-b). Overall, the Fourier Shell Correlation (FSC) resolution of the reconstructed maps (half-maps at threshold 0.143) was measured to be 2.8 Å for the Compound B1 dataset and 3.2 Å for the Compound B2 dataset (Fig. 6 a-b) compared to 2.4 Å and 2.6 Å obtained with a standard workflow of 2D classification and auto-refine in RELION, but at a fraction of the computational cost (see Results Section 2.4). Similarly, the map-to-model FSC resolutions (threshold 0.5) were 2.9 Å and 3.2 Å, confirming good agreement between the reconstructed maps and their corresponding ground-truth atomic models.

**Fig. 2.**
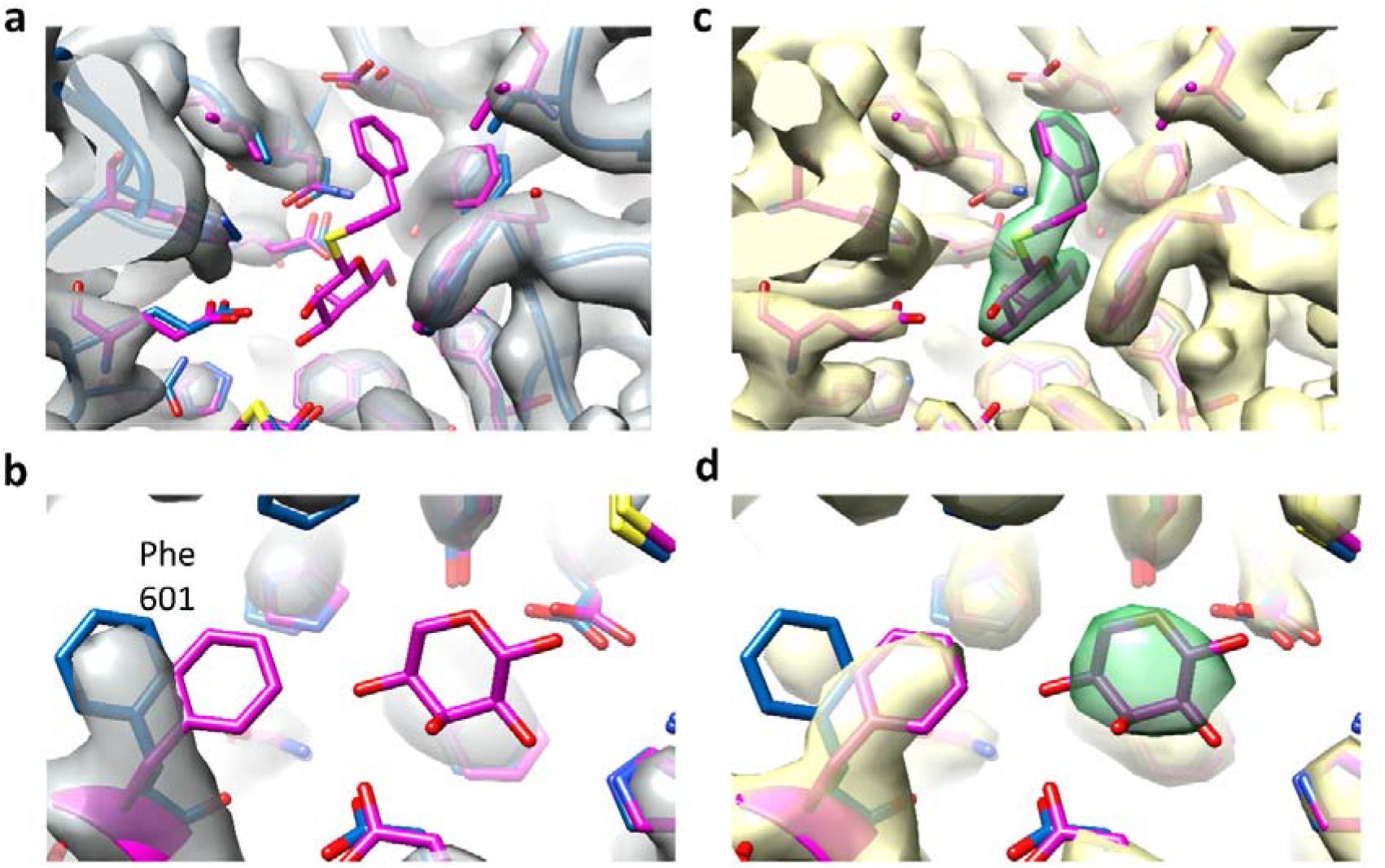
Reconstructed maps of β-galactosidase (BGAL) at the ligand binding site using orientations inferred with cryoPARES. a) Map of the apoprotein (grey) with its associated atomic model (blue) and the aligned atomic model of the protein bound to Compound B1 (pink). b) Same as (a) but with the atomic model of the protein bound to Compound B2 (pink). c) Reconstructed map (yellow) of the protein bound to Compound B1 using orientations estimated with cryoPARES. The ligand and residues within 4 Å of the ligand are displayed in pink. The density closer than 1.8 Å to the ligand is coloured in green. d) Same as (c), but for the protein bound to Compound B2. The features for the ligands are absent in the apo maps (a, b) but clearly visible in the reconstructed maps from the bound datasets (c, d). In this case, the binding of ligands does not induce significant changes in the structure of the protein pocket, with the exception of the Phe 601 in the Compound B2 case (b vs d).

Our second test was conducted on our previously published datasets of human pyruvate kinase M2 (PKM2) bound to Compounds P1 and P2^3^. As with BGAL, the densities for the ligand-bound reconstructions obtained using cryoPARES poses matched the ground-truth atomic models (Fig. 3). In this case, the FSC resolution obtained with cryoPARES poses was closer to the resolution obtained with the standard RELION workflow: 3.5 Å and 3.3 Å for cryoPARES (Fig. 6 c-d), compared to 3.4 Å and 2.8 Å. The map-to-model resolutions were 3.7 Å and 3.5 Å. For this protein target, the apo dataset used for training was collected on a Krios microscope, but the PKM2 Compound P1 dataset was collected on a Glacios microscope, demonstrating the method is robust under different imaging conditions.

**Fig. 3.**
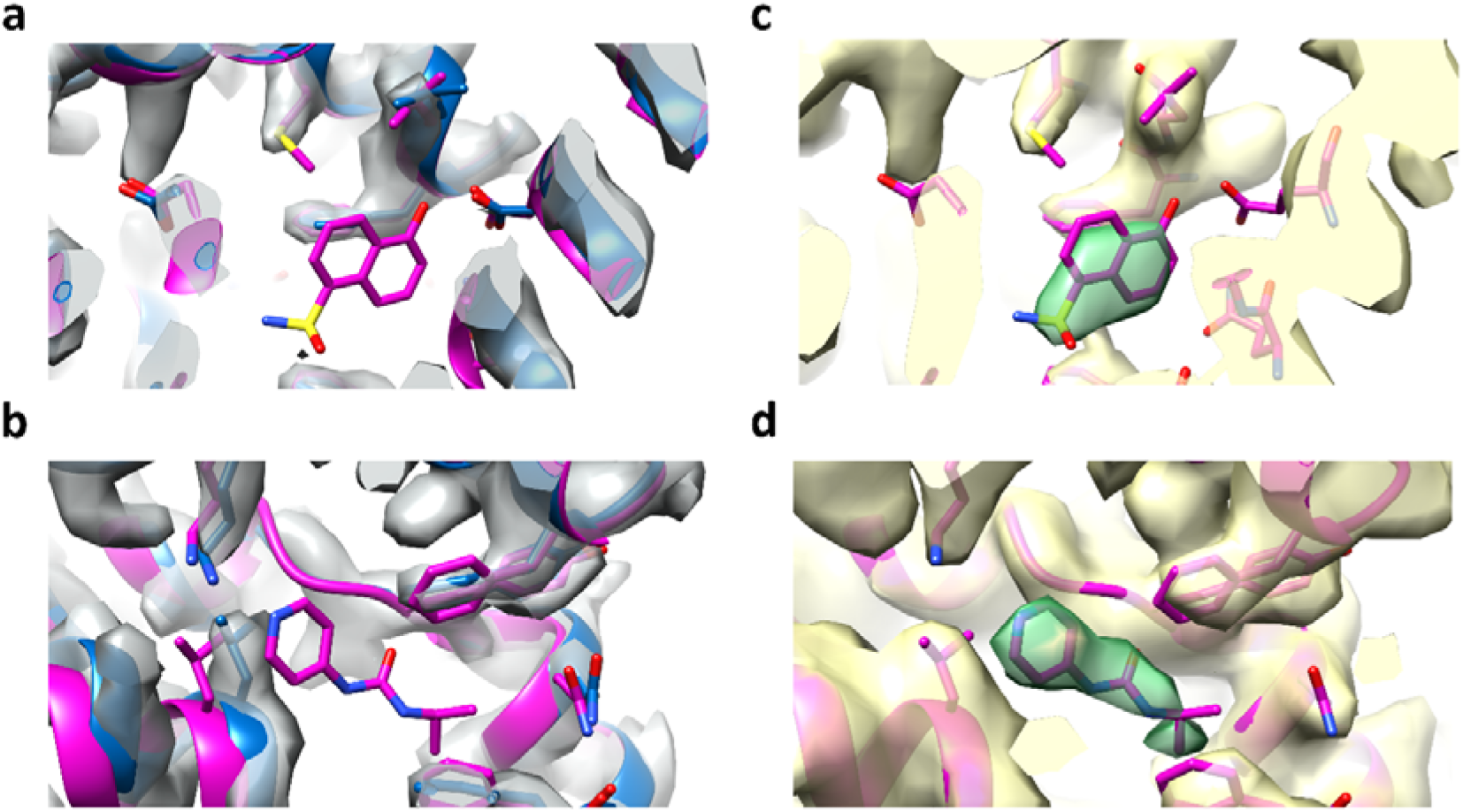
Reconstructed maps of pyruvate kinase M2 (PKM2) at the ligand binding site using orientations inferred with cryoPARES. a) Density of the apoprotein (grey) with its associated atomic model (blue) and the aligned atomic model of the protein bound to Compound P1 (pink). b) Same as (a) but with the atomic model of the protein bound to Compound P2 (pink). c) Reconstructed density (yellow) of the protein bound to Compound P1 using orientations estimated with cryoPARES. The ligand and residues within 4 Å of the ligand are displayed in pink. The density closer than 1.8 Å to the ligand is coloured in green. d) Same as (c), but for the protein bound to Compound P2. The density for the ligands is absent in the apo maps (a, b) but clearly visible in the reconstructed maps from the bound datasets (c, d).

The third case consists of two in-house datasets of bovine glutamate dehydrogenase (GDH) bound to Compounds G1 and G2. Fig. 4 shows the maps for the apo structure used in the training and local refinement steps, as well as the densities of the maps reconstructed from cryoPARES-predicted poses. In this case, we measured FSC resolutions of 2.9 Å, and 3.0 Å (see Fig. 6 e-f), compared to 2.7 Å and 2.8 Å for the RELION workflow. The map-to-model resolutions were 2.9 Å and 3.1 Å.

**Fig. 4.**
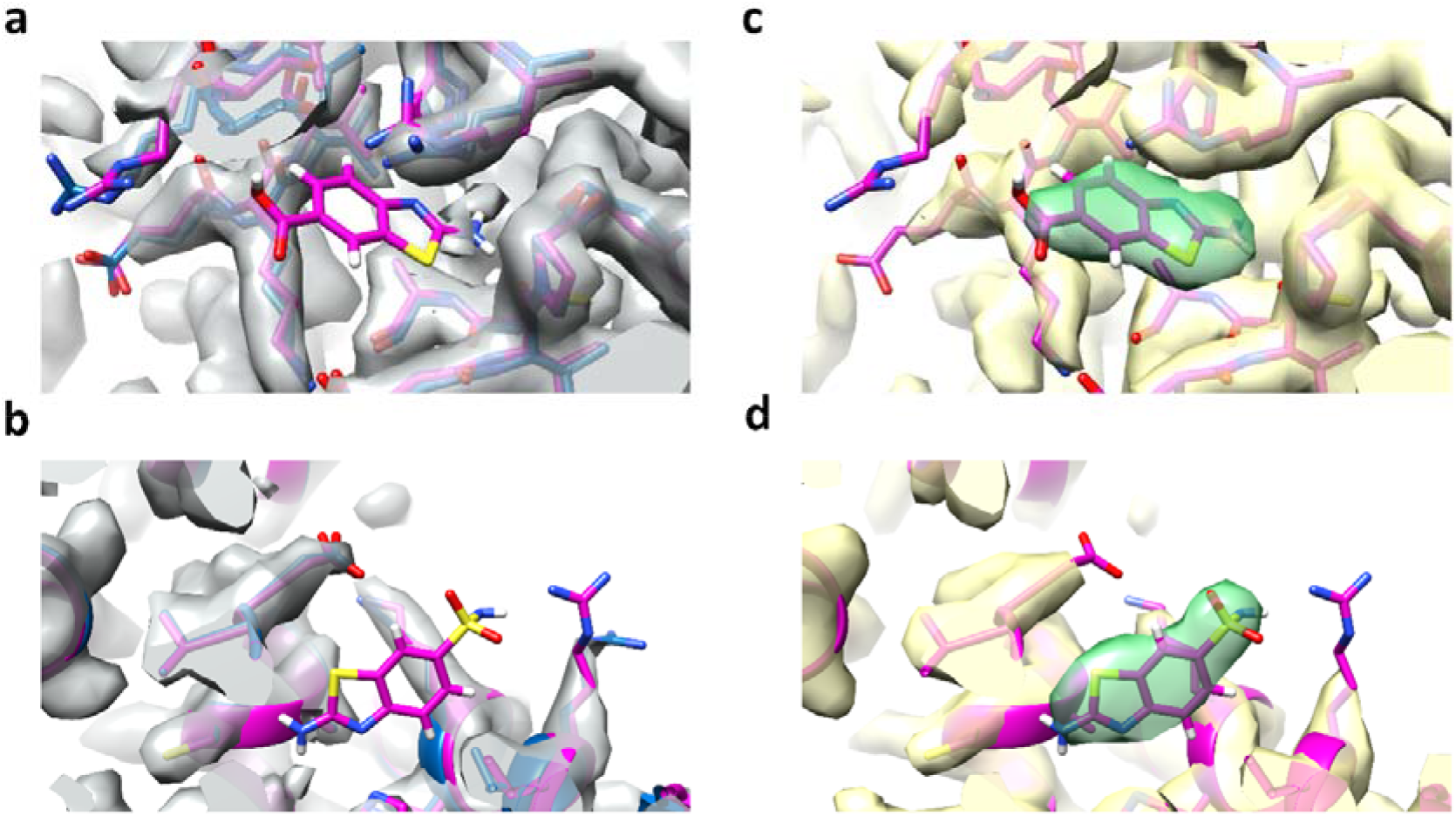
Reconstructed maps of glutamate dehydrogenase (GDH) at the ligand binding site using orientations inferred with cryoPARES. a) Density of the apoprotein (grey) with its associated atomic model (blue) and the aligned atomic model of the protein bound to Compound G1 (pink). b) Same as (a) but with the atomic model of the protein bound to Compound G2 (pink). c) Reconstructed density (yellow) of the protein bound to Compound G1 using orientations estimated with cryoPARES. The ligand and residues within 4 Å of the ligand are displayed in pink. The density closer than 1.8 Å to the ligand is coloured in green. d) Same as (c), but for the protein bound to Compound G2. The density for the ligands is absent in the apo maps (a, b) but clearly visible in the reconstructed maps from the bound datasets (c, d).

The final test case consists of an in-house dataset of transient receptor potential mucolipin 1 (TRPML1) bound to Compound T1^29^. Fig. 5 shows the map for the apo structure of this ion channel used in the training and local refinement steps, as well as the density of the map reconstructed from cryoPARES-predicted poses. In this case, we measured an FSC resolution of 2.7 Å (see Fig. 6 g), compared to 2.1 Å for a rather complex cryoSPARC^7^ workflow described in Reeks et al.^29^. The map-to-model resolution for the cryoPARES map was 2.9 Å. For an alternative visualization of the ligands shown in Figures 2-5, see Supplementary Fig. 8.

**Fig. 5.**
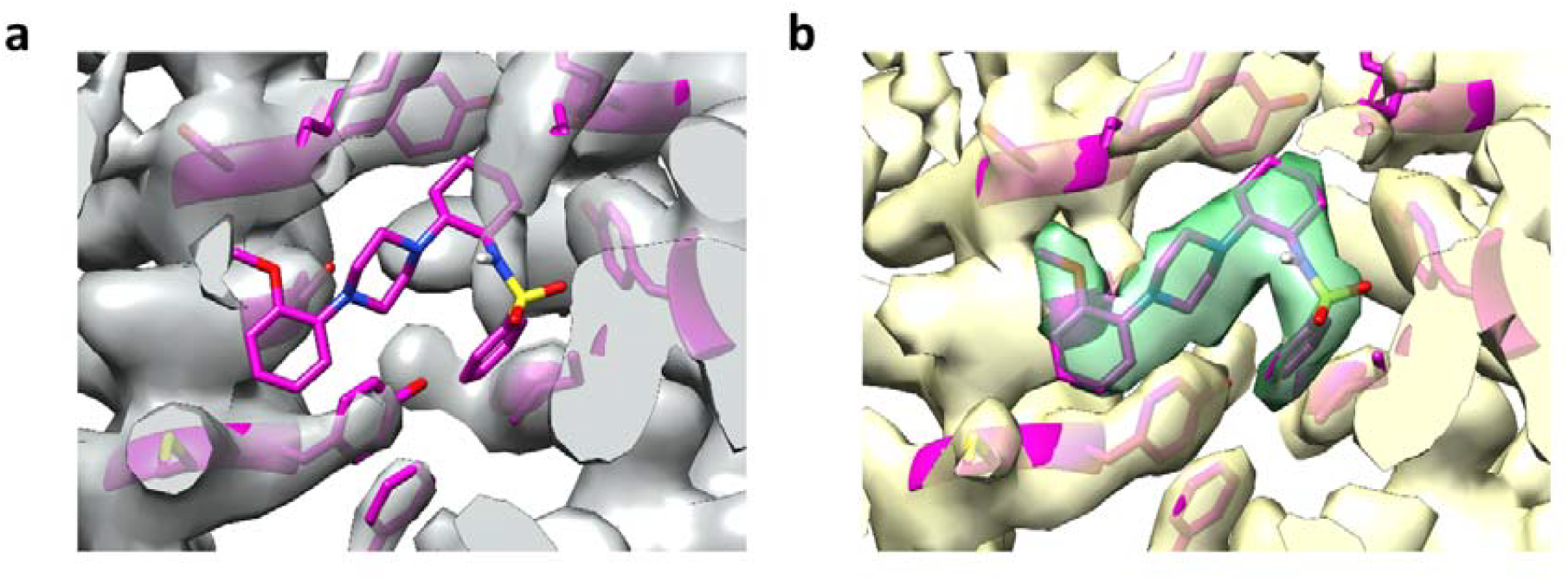
Reconstructed maps of human transient receptor potential mucolipin 1 (TRPML1) at the ligand binding site using orientations inferred with cryoPARES. a) Density of the apoprotein (grey) and the aligned atomic model of the protein bound to Compound T1 (pink). b) Reconstructed density (yellow) of the protein bound to Compound T1 using orientations estimated with cryoPARES and the aligned atomic model of the protein bound to Compound T1 (pink). Only the ligand and residues within 4 Å of the ligand are displayed. The density closer than 1.8 Å to the ligand is coloured in green. The density for the ligand is absent in the apo map (a) but clearly visible in the reconstructed map from the bound datasets (b).

These four experiments confirm that cryoPARES can deliver high-resolution structures of ligands, (in the 3.5 to <3.0 Å resolution range, see Supplementary Fig. 9 for local resolution estimations). While FSC measurements can be affected by template bias, there are several observations strongly indicating that the effect of this, if present, is small: 1) the ligands reconstructed were neither present in the training data nor the template, 2) the measured resolutions of ∼3 Å are significantly better than the frequency threshold used to low-pass filter the local refinement templates (6 Å), and 3) the density of phenylalanine 601 in Fig. 2 b,d shows a clear, known rotamer change upon ligand binding. Further tests and discussion on template bias avoidance and detection can be found in Supplementary Note 7, Supplementary Table 9, and Supplementary Fig. 6.

Furthermore, while the ligand-binding cases presented in this section focus on relatively large and symmetric proteins, we also evaluated our method using the CESPED benchmark (Supplementary Note 8). In these experiments, a half-dataset of particles is used to train the model, which is then used to infer the poses of the remaining half-dataset. Results in Supplementary Table 10 confirm that the method remains robust for smaller targets (<200 kDa) and asymmetric systems, demonstrating its broader applicability.

Finally, we briefly address sensitivity of cryoPARES to conformational changes, i.e. conformational differences between the apo data used for training and the ligand-bound samples used for inference. Based on our experiments, moderate structural deviations appear to be well-tolerated, as demonstrated by the successful reconstruction of PKM2 despite a backbone RMSD of 2.1 Å and 2.7 Å for P1 and P2 (Supplementary Note 9 and Supplementary Table 12). Notably, the P1 state of PKM2 exhibited a lower RMSD than P2 yet resulted in a lower resolution reconstruction, suggesting that structural divergence at these levels is not the primary driver of performance loss.

### 2.3. CryoPARES predicts accurate global alignment parameters

We first evaluated accuracy on simulated data with perfect orientations, where the cryoPARES’s neural network achieved validation top-1 median angular errors of 3.9°, 4.6°, and 4.1° for BGAL, PKM2 and GDH protein targets, approaching the SO(3) grid resolution of ∼3.7° used within the model. While these results demonstrate the model’s capability under ideal conditions, evaluating angular errors on real datasets is more challenging due to the lack of ground truth poses.

For experimental datasets, we compared our predicted poses with those from RELION (see Supplementary Table 4). The neural network’s top-1 raw predictions are within 5° of RELION estimates for approximately 30% of particles, increasing to more than 60% when followed by local refinement. Notably, even two consecutive RELION auto-refine jobs on the same dataset can differ by more than 5° for over 20% of particles (see Supplementary Table 4, last column), with studies reporting disagreement levels as high as 50%^23^. This substantial variability between identical RELION runs underscores that perfect agreement between methods is neither expected nor necessarily desirable, since it could reflect shared systematic bias rather than true accuracy. Therefore, our observed level of correspondence with RELION, combined with the quality of our reconstructed maps, supports the reliability of cryoPARES poses.

Another indirect method for assessing the accuracy of the neural network angular assignment is to measure how much local refinement is needed to improve the reconstruction before no gains are observed. Fig. 6 shows the FSC curves for all the samples where local refinement with an angular search range of 0°, ±2°, ±4° and ±6° and step size of 2° was applied. In every dataset, most of the gain was achieved with a ± 2° search, with only marginal improvement beyond ± 4°, indicating that the predicted orientations are typically within 2–4° of the optimal alignment.

**Fig. 6.**
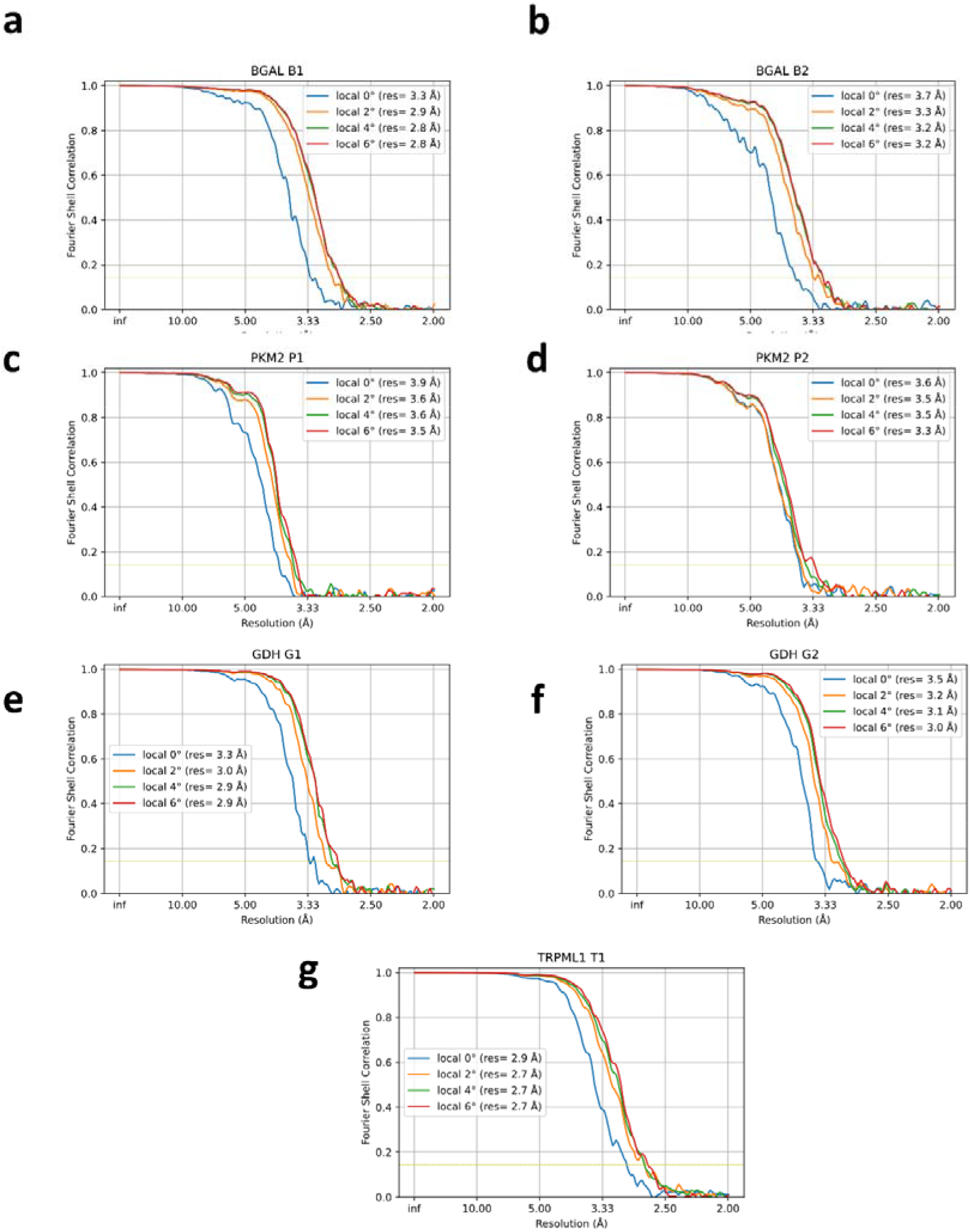
FSC curves (half-maps, 0.143 resolution threshold) for the maps obtained by cryoPARES at different levels of local refinement angular search range (0° blue, ±2° orange, ±4° green, ±6° red) for the protein-ligand complexes BGAL bound to Compound B1 (a) and Compound B2 (b), PKM2 bound to Compound P1 (c) and Compound P2 (d), GDH bound to Compound G1 (e) and Compound G2 (f), TRPML1 bound to Compound T1 (g). The FSC resolution at 0.143 is reported in parentheses.

While these metrics confirm the reliability of the global pose assignments, the network’s performance scales further with data availability (Supplementary Table 14). These results indicate that the initial neural network predictions are sufficiently accurate for high-resolution convergence and are not the primary cause of the slight resolution gap observed relative to RELION. Instead, this difference stems from our local refinement implementation, which prioritizes speed for real-time screening. To bridge this gap, we have implemented an optional, more complex refinement algorithm that utilizes two projection matching stages together with interpolation strategies to more precisely locate cross-correlation peaks in the SO(3) and pixel grids. While this enhanced protocol is 2–3× slower, it provides significant improvements in both angular accuracy and final FSC resolution, as detailed in Supplementary Table 15. This advanced mode allows cryoPARES to achieve results much closer to traditional pipelines while maintaining its automated, on-the-fly nature.

For a comparison of cryoPARES accuracy with similar approaches, see Supplementary Note 8 and Supplementary Table 11. A sensitivity analysis with respect to training set quality is provided in Supplementary Note 10 and Supplementary Table 13.

### 2.4. CryoPARES can perform real-time particle alignment

Supplementary Table 6 summarizes the measured runtimes for pose estimation across the different datasets. During inference, the cryoPARES neural network achieves throughputs exceeding 30,000 particles min^−1^ GPU^−1^, allowing a 250,000-particle stack to be processed in less than eight minutes. If we assume that cryoPARES rejects as junk particles approximately 50% of the dataset, the local refinement of the remaining particles would take about seven additional minutes, resulting in a total runtime of roughly 15 minutes on a single GPU for 250,000 picked particles.

Since our local refinement implementation is a proof-of-concept demonstrator, performance could be further enhanced through better alignment algorithms or by reducing the amount of local refinement required via improvements of the neural network. Even in its present form, cryoPARES is more than an order of magnitude faster than RELION auto-refinement and comparable in speed to the non-open source cryoSPARC homogeneous refinement (see Supplementary Table 7). Given that a high-end microscope can collect 700+ micrographs h^-1^, and assuming 400 particles per micrograph, real-time processing requires ∼4,700 particles min^−1^. CryoPARES comfortably exceeds even the most demanding acquisition rates, sustaining > 10,000 particles min^−1^ GPU^−1^ including the local-refinement step.

Unlike conventional workflows that require accumulating a minimal number of particles to perform statistical estimations and preprocessing steps like 2D classification, cryoPARES processes each particle independently. This fundamental difference not only enables processing to begin immediately after the first micrograph is acquired but also allows for continuous updates of the 3D reconstruction as new particles become available. Together, the combination of high-throughput processing and per-particle analysis has the potential to transform cryo-EM data processing from a batch-based, post-acquisition task into a real-time monitoring system that provides immediate feedback on data quality and experimental parameters.

### 2.5. CryoPARES can perform automatic particle pruning

CryoPARES provides probability distributions over particle orientations, which can be used to identify junk particles and those with unreliable estimated poses. While one could directly use the probability of the predicted orientation as a quality metric, we observed that the distribution of these raw probabilities depends strongly on viewing direction. Applying a single global threshold would therefore bias reconstructions toward more frequent orientations, potentially producing artefactual preferred orientations (Supplementary Fig. 3).

To overcome this, we implemented a pruning score based on direction-normalized robust z-scores of the predicted probability distributions (see Methods Section 4.10). This normalization allows particle and pose quality to be evaluated consistently across all orientations. Supplementary Fig. 4 presents histograms of the direction-normalized scores for “good” particles (those retained after 2D classification) and “bad” particles (those discarded). In every case, the distributions for good particles extend toward higher scores, whereas those for bad particles are concentrated at lower values, confirming that a single threshold can separate the two populations. All reconstructions reported in the previous sections were obtained from the datasets pruned using automatically estimated thresholds of this score (see Supplementary Note 3 and Supplementary Fig. 2).

To further assess pruning performance, we compared RELION auto-refine results obtained using (i) particles cleaned by 2D classification, (ii) particles selected using cryoPARES pruning scores, and (iii) randomly chosen subsets containing the same number of particles. Note that, for this experiment only, we manually adjusted the cryoPARES thresholds to match the number of retained particles from 2D classification. For this reason, the thresholds used here differ from the automatically determined values employed in the previous sections. Table 1 summarizes the resulting resolutions. Datasets pruned by cryoPARES achieved resolutions comparable to those obtained through manual 2D classification, while random subsets consistently produced lower-resolution reconstructions. Because the datasets analysed so far were relatively clean (as evidenced by the small effect of random subsetting in several cases), we also evaluated cryoPARES pruning on two in-house TRPML1 liganded dataset, different from T1 (TRPML1 α and TRPML1 β), which contain a higher proportion of poorly aligned and junk particles. In these more challenging cases, the benefits of pruning were particularly pronounced: cryoPARES-selected particles improved the final resolutions by 0.4–1.2 Å relative to random subsets and reached values nearly identical to those obtained after exhaustive 2D classification.

**Table 1.** Resolution obtained with a RELION auto-refine job when the input particles are (1) cleaned with a 2D classification step, (2) cryoPARES directional pruning scores, or (3) randomly selected particles. The same number of particles in the second and third cases was selected to match the 2D classification outcome.

| Sample | Resolution after 2D classification (Å) | Resolution after cryoPARES (Å) | Resolution after random subset (Å) |
| --- | --- | --- | --- |
| BGAL Compound B1 | 2.51 | 2.50 | 2.74 |
| BGAL Compound B2 | 2.75 | 2.70 | 3.10 |
| PKM2 Compound P1 | 3.32 | 3.36 | 3.39 |
| PKM2 Compound P2 | 2.88 | 2.82 | 2.96 |
| GDH Compound G1 | 2.52 | 2.50 | 2.54 |
| GDH Compound G2 | 2.67 | 2.63 | 2.74 |
| TRPML1 T1 | 3.28 | 2.92 | 3.52 |
| TRPML1 $\alpha$ | 2.54 | 2.57 | 3.02 |
| TRPML1 $\beta$ | 2.79 | 2.67 | 3.92 |

Additionally, we conducted two experiments using two types of out-of-distribution (OOD) inputs: pure Gaussian noise and a cross-contamination dataset created by mixing particles from two unrelated protein systems (BGAL and TRPML1). As shown in Fig. 7, cryoPARES assigned significantly lower pruning scores to the OOD inputs compared to the in-distribution particles in both scenarios, effectively preventing noise and mismatched data from contributing to the final reconstruction. When interpreting these distributions, it is important to note that the pruning score functions as a measure of alignment confidence rather than a strict binary “junk” detector. Consequently, any input that cannot be reliably aligned—whether it is a “junk” particle or a genuine particle with a poor signal-to-noise ratio—is assigned a lower score. This can explain why the distribution of in-distribution particles is notably wider than that of the OOD inputs. For a characterization of the pruned particles, see Supplementary Note 6 and Supplementary Fig. 5.

**Fig. 7.**
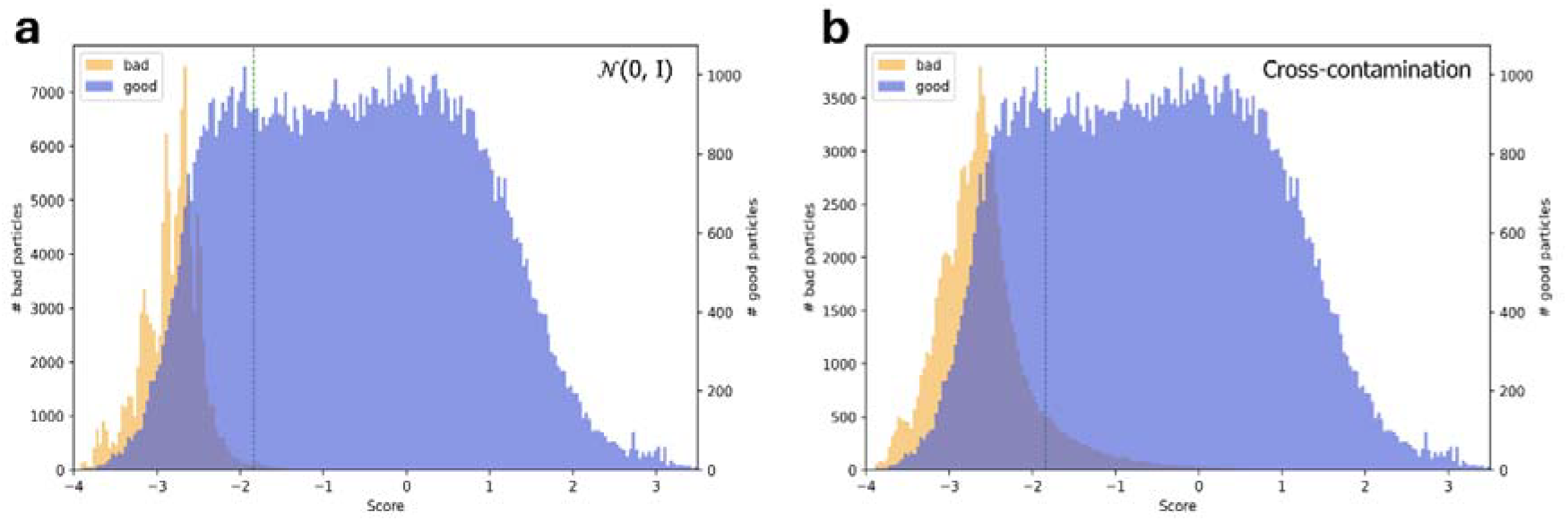
Distribution of direction-normalized pruning scores for out-of-distribution inputs compared to in-distribution particles. Histograms show pruning scores computed with a model trained on TRPML1 apo particles for (a) pure Gaussian noise particles (*N* (0,1) and (b) particles from BGAL, an unrelated protein (Cross-contamination) (orange), compared to TRPML1 T1 particles (blue). In both cases, out-of-distribution particles are shifted towards lower pruning scores. Note that the pruning score reflects alignment reliability rather than a strict junk/good classification, which can explain the partial overlap between distributions. Different y-axis scales are used due to the imbalance in the number of in-distribution and out-of-distribution particles. The vertical green line corresponds to the threshold automatically estimated. The shadowed region corresponds to overlapping bins.

Together, these analyses show that cryoPARES can effectively replace 2D classification as a particle-pruning step, offering two key advantages: on-the-fly operation and full reproducibility, as no manual class selection is required.

### 2.6. CryoPARES shows potential for alleviating preferred orientation bias

CryoPARES, similar to baited reconstruction approaches^19^ avoids the attractor effect that arises in iterative refinement. It happens when orientations represented by more particles—and thus higher signal-to-noise ratios—pull neighbouring particles toward themselves, amplifying over-represented directions and biasing the reconstruction^30,31^. This property suggests that cryoPARES should be less susceptible to preferred orientations than iterative refinement methods, since its training on unbiased pose distributions prevents reinforcement of dominant views.

To illustrate this situation, we analysed another GDH dataset (bound to ADP, Compound G3) exhibiting significant preferred orientations. When reconstructed using a standard RELION workflow —comprising 2D classification followed by auto-refine with a 40 Å low-pass filtered reference— the resulting volume shows severe anisotropy that precludes ligand identification (Fig. 8 a). However, employing a higher-resolution density (15 Å) as reference yields substantially improved reconstruction quality, enabling ligand visualization (Fig. 8 b,d). This scenario resembles the baited reconstruction strategy, where a high-resolution reference is used as template.

**Fig. 8.**
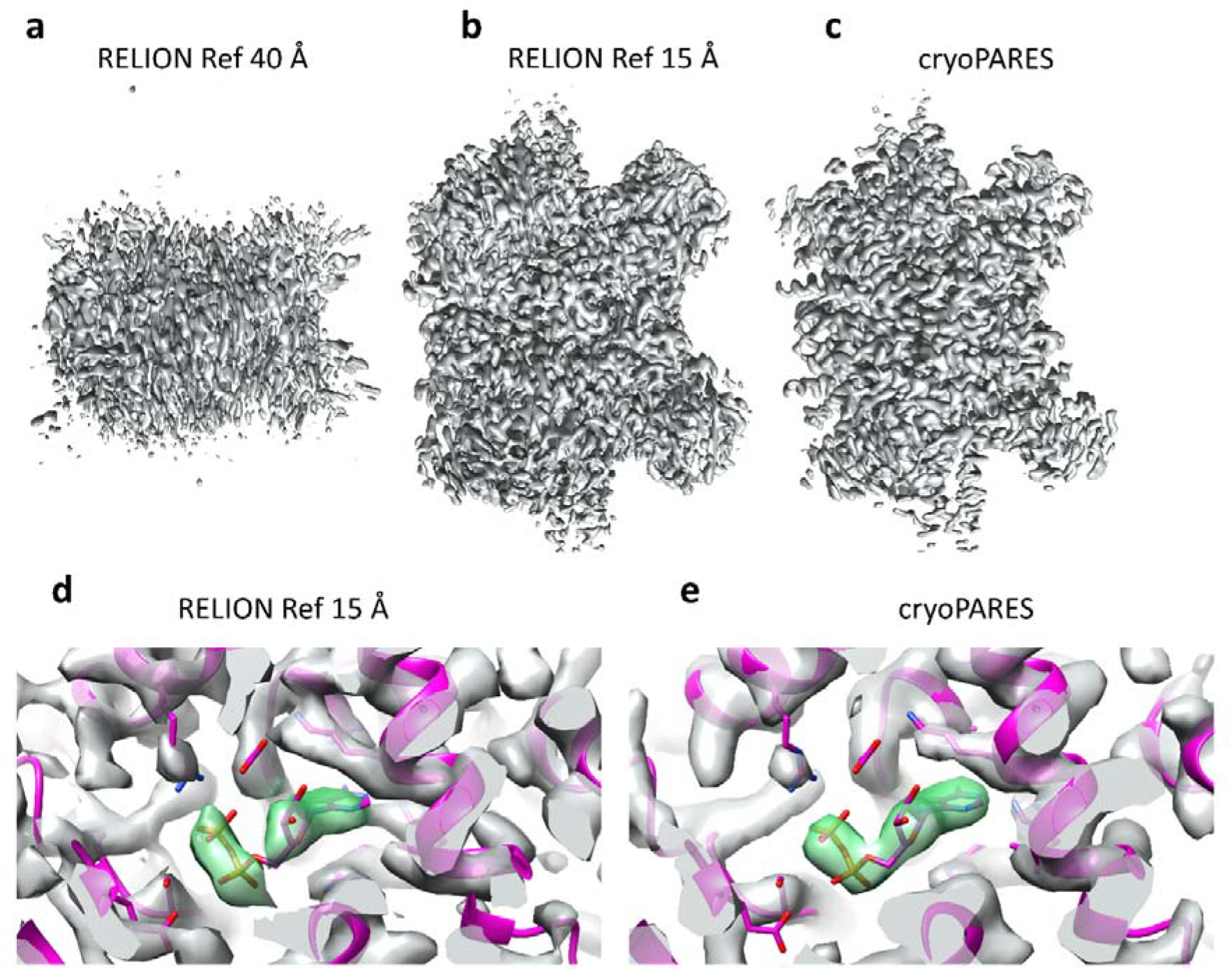
Reconstructed densities from a dataset of GDH bound to ADP that contains preferred orientations. a) Overview of the map obtained with a RELION workflow of 2D classification followed by auto-refine with a reference low-pass filtered at 40 Å b) Same as (a) with a reference low-pass filtered at 15 Å. c) Overview of the map reconstructed using cryoPARES. d) Close-up of (b) showing the ADP binding site. e) Close-up of (c) showing the ADP binding site. The atomic model (pink) corresponds to the ground-truth structure, computed from another stack of ADP-bound GDH particles not affected by preferred orientations. In panels d-e, the density within 1.8 Å of the ADP is coloured in green.

By utilizing the same cryoPARES model as before—trained on the apo-dataset, unaffected by preferred orientation—we were able to reconstruct a map with correct global features and markedly better-resolved local details (Fig. 8 c,e) and improved anisotropy statistics (see Supplementary Fig. 7). Notably, the ligand density in the cryoPARES map was more continuous than in the RELION result, and nearby side-chains—such as LYS488, adjacent to the ADP molecule—exhibited significantly better definition. These results confirm that, for this challenging dataset, cryoPARES substantially alleviates the effects of preferred orientation. While additional datasets will be required to establish the generality of this behaviour, these findings are consistent with the expectation that the absence of the attractor effect in CryoPARES can mitigate reconstruction bias caused by preferred orientations.

## 3. Discussion

In this work, we introduced cryoPARES, a supervised deep-learning framework that performs efficient cryo-EM pose determination in cases where pre-aligned particle datasets of related samples are available. By learning from these datasets, cryoPARES rapidly predicts particle orientations while performing automatic particle pruning, thereby significantly reducing the computational burden of single-particle analysis.

Experiments with BGAL, PKM2, GDH and TRPML1 demonstrate that cryoPARES reconstructs high-resolution ligand-bound complexes that accurately reproduce ligand densities absent in the apo structures used for training. The method generalizes across different datasets and imaging conditions, maintaining high performance even when trained on data acquired with multiple microscopes. These results support its applicability to realistic drug discovery workflows. In addition, our results indicate that cryoPARES has the potential to alleviate limitations associated with iterative refinement, such as sensitivity to preferred orientations. While this observation is currently supported by a single experimental dataset, it is consistent with the absence of the attractor effect in amortized pose inference and motivates further investigation across a broader range of samples.

The high-speed, one-shot nature of cryoPARES makes real-time data analysis during cryo-EM acquisition feasible, providing immediate feedback on data quality and experimental parameters. This capability, combined with automatic particle pruning, is particularly valuable for high-throughput structural studies and drug discovery campaigns, where rapid iteration, automation, and reproducibility are essential.

The present study also highlights several directions for future work. Although the current implementation already achieves reconstructions close to those obtained with conventional iterative pipelines, further improvements in pose prediction and local refinement could narrow the remaining resolution gap without sacrificing real-time performance. More broadly, extending the supervised amortized inference paradigm beyond closely related samples, accounting for conformational variability, and evaluating performance on a wider range of challenging datasets with preferred orientations represent promising directions for future research. Together, these results establish cryoPARES as a practical framework for accelerating cryo-EM studies in structure-based drug discovery and highlight the broader potential of supervised pose inference for high-throughput structural biology.

## 4. Methods

### 4.1. Datasets

We studied four protein targets, β-galactosidase (BGAL), pyruvate kinase M2 (PKM2), glutamate dehydrogenase (GDH) and human transient receptor potential mucolipin 1 (TRPML1). For each protein target, we prepared two types of datasets: training data from apo samples and testing data from ligand-bound samples (see Supplementary Table 2 for a list of compounds).

The testing sets for the BGAL, PKM2 and TRPML1 targets were prepared from previous work^3,29^: EMPIAR-10644 (β-galactosidase bound to PETG, Compound B1), EMPIAR-10646 (β-galactosidase bound to L-ribose, Compound B2), EMPIAR-10648 (PKM2 bound to 5-hydroxynaphthalene-1-sulfonamide, Compound P1), EMPIAR-10649 (PKM2 bound to 3-(propan-2-yl)-1-(pyridin-4-yl)urea, Compound P2), and EMDB-52249 (TRPML1 bound to N-[2-[4-(2-methoxyphenyl)piperazin-1-yl]phenyl]benzenesulfonamide). The training sets for BGAL, PKM2 and TRPML1 were prepared following an identical protocol as the ligand-bound datasets.

Please note that the training set for TRPML1, as well as the ground-truth angles and reported resolution for TRPML1 T1, were obtained using a more complex cryoSPARC workflow (described in ^29^) rather than the simpler RELION workflow used for the other targets. In contrast, the results presented in Results Section 2.5 for TRPML1 were obtained using a single RELION 2D classification step followed by a RELION auto-refine step, consistent with the other examples.

For GDH, the employed ligands were 2-amino-1,3-benzothiazole-6-carboxylic acid (Compound G1), and 2-amino-1,3-benzothiazole-6-sulfonamide (Compound G2). An additional dataset, ADP-bound (Compound G3), was employed for the preferred orientations analysis.

### 4.2. Glutamate dehydrogenase (GDH) protein preparation and cryo-EM data collection

Bovine glutamate dehydrogenase (GDH) was obtained from Sigma-Aldrich as a 17⍰mg/mL solution. It was dialyzed overnight at 4 °C against 100⍰mM HEPES, 150 mM NaCl, pH 8.0. The protein was further purified using size-exclusion chromatography on a HiLoad 16/60 Superdex™ 200⍰pg column (Cytiva) using the same buffer. Fractions containing GDH were pooled and concentrated using Ultracel 10K centrifugal filters (Amicon) to a final concentration of 2.35⍰mg/mL. The protein was snap frozen in liquid nitrogen and stored at -80 °C until required. For the protein sample used for compound G3 only, the buffer was instead 100 mM sodium phosphate, pH 6.8 and the final protein concentration was 3.5⍰mg/mL.

Samples for cryo-EM were prepared as follows. Apo: GDH was diluted to 0.35 mg/mL in 100 mM HEPES, 150 mM NaCl, pH 8.0 supplemented with 5 % DMSO and 0.001 % Tween-20. Compounds G1 and G2: compound was dissolved as a 2 M stock in DMSO. It was then diluted to 120 mM, 6 % DMSO in 200 mM HEPES, 150 mM NaCl, pH 8.0 and then the pH was manually re-adjusted to pH 8.0. The compound was then mixed with GDH and Tween-20 to give final concentrations of 100 mM compound in 5 % DMSO, 0.35 mg/mL GDH and 0.001 % Tween-20. Compound G3: compound was dissolved as a 100 mM stock in 100 mM sodium phosphate, pH 6.8. The compound was then mixed with DMSO, GDH, and Tween-20 to give final concentrations of 20 mM compound with 5 % DMSO, 0.35 mg/mL GDH and 0.001 % Tween-20. Protein-ligand complexes were incubated on ice for 30 minutes before grid preparation.

Graphene oxide (Sigma-Aldrich) was applied to R1.2/1.3 300 mesh holey carbon grids (Quantifoil Micro Tools) as described by Martin T.G. et al.^32^. 3 μL GDH sample was applied to a grid and blotted using a Vitrobot Mark IV (Thermo Fisher Scientific) at 4 °C and 100 % relative humidity before being plunged into liquid ethane.

Cryo-EM data were collected on a Thermo Fisher Scientific Titan Krios TEM at the Cambridge Pharmaceutical Consortium (apo and Compound G1 and G2) and the Astbury BioStructure Laboratory (Compound G3). Micrographs were recorded using a Falcon3 direct electron detector in counting mode. Supplementary Table 1 lists important experimental conditions and results.

### 4.3. Glutamate dehydrogenase (GDH) atomic modelling

Molecular replacement was used to fit the reference atomic model into the cryo-EM maps. To do so, structure factors were calculated from the cryo-EM density maps, and each chain of the assembly was independently positioned using Phaser^33^ version 2.8.2. Initial refinement was performed using Coot min-rsr 0.9.6^35^ to optimize the fit between the protein structure and the experimental density, primarily involving minor adjustments to side chain orientations. Where necessary, larger conformational changes in the protein backbone were manually adjusted using WinCoot^35^ version 0.9.8.96. Once the protein structure adequately accounted for all observed density features, the remaining ligand density was interpreted and fitted using the automated procedures in AutoSolve™^36^ build number 230907, followed by a real-space refinement in Phenix 1.21rc1^37^.

### 4.4. Datasets image processing

The particle extraction process was identical for both training and testing datasets. CTF estimation was performed using Gctf 1.18^38^ (BGAL, PKM2, GDH) or cryoSPARC^7^ Patch CTF (TRPML1), using a 1024-pixel box size, 30 Å minimum resolution, and 0.07 amplitude contrast. We then conducted particle picking via template matching using Gautomatch 0.56 (http://www.mrc-lmb.cam.ac.uk/kzhang/, BGAL, PKM2, GDH) or cryoSPARC v4.7.1 template picker (TRPML1), with references low-pass filtered to 20 Å, and extracted all particles at 1 Å/pixel. For testing, these extracted particles were directly fed to cryoPARES, which predicts their poses without requiring any additional processing. However, to train cryoPARES and to evaluate cryoPARES predictions, we needed reference poses and an estimation of good and bad particles for particle pruning. We obtained these by processing both training and testing datasets through a 2D classification job with 25 steps in RELION 3^39^, followed by 3D refinement using RELION auto-refine with a reference map low-pass filtered at 20 Å (BGAL and PKM2, TRPML1) or 15 Å (GDH) and a soft mask created from the reference map. See Supplementary Table 1 and Supplementary Fig. 10-12 for additional details.

### 4.5. Training strategy

For each of the targets, the apo dataset containing only “good” particles is evenly split into two half-sets of particles. Then, for each of the half-sets, a model is trained using 70% of the data as the training set and the remaining 30% as the validation set. In each of the training runs, we first pre-train our model for 5 epochs using a simulated dataset of particles (see Supplementary Fig. 1 for more details about the simulation). After that, we continue training for at most 100 epochs with the training set of experimental particles. While the pre-training phase using simulated data is not necessary – as models can be directly trained on experimental particles alone – it can lead to better results in some cases (see Supplementary Fig. 1).

For both the pre-training and the training steps, we employ RAdam^40^ as the optimizer, with an initial learning rate of 1e-3 and a weight decay of 5e-6. The metric used to monitor the training progress was the mean validation geodesic distance,

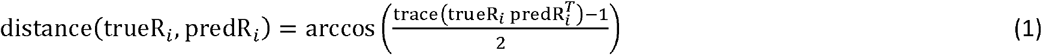

where trueR_*i*_ and predR_*i*_ are, respectively, the ground-truth and predicted rotation matrix for the *i*- particle of the validation set. When not stated otherwise, predR_i_ refers to the highest probability rotation matrix, referred as the top-1 orientation. During training, we halve the learning rate if the validation error did not improve for more than 5 epochs. Early stopping is applied when the validation error did not improve for more than 11 epochs. Data augmentation is applied for the training set (see Supplementary Note 1 for details). We use a batch size of 32 images with gradient accumulation over 20 batches. Each batch includes 4 copies of the same particles subjected to different data augmentation techniques. This is intended to help the network learn augmentation-invariant features, particularly for shifts and in-plane rotations. An optional SimCLR-like contrastive loss^41^ was also implemented to favour learning such features. However, no significant performance gains for the datasets presented here were observed.

Once training is completed, we apply the trained model to predict poses for particles in the validation set. When comparing these predicted poses with their ground truth values, we also compute the metrics needed to develop our particle pruning criteria based on direction-normalized robust z-scores, which help us filter out junk and/or unreliably aligned particles in new datasets (see Methods Section 4.10 for details).

### 4.6. Pose inference strategy

For pose inference, we start with the ligand-bound particles extracted directly after particle picking. We split these particles into two equally-sized subsets and assign each subset to one of our two independently trained apoprotein models for initial pose prediction. Following this initial prediction, we employ our particle pruning strategy based on direction-normalized robust z-scores to filter out junk particles and incorrectly aligned particles (see Methods Section 4.10). The remaining particles then undergo local refinement to improve their orientations and/or estimate in-plane shifts (see Methods Section 4.9).

### 4.7. Data pre-processing

Before being fed to the neural network, the particle images undergo several preprocessing steps. First, they are downsampled to 1.5 Å/pixel and normalized such that the corner regions, which should contain only noise, exhibit a mean of 0 and standard deviation of 1. The images are then enriched with an additional channel containing their CTF-corrected (phase-flipped) versions. From these processed images, we extract central crops. Then, a circular mask is applied with a diameter equal to the particle diameter. To enhance feature detection across multiple levels of noise, as suggested by Levy et al.^13^, we apply a series of Gaussian filters (σ = 0, 0.3, 1, and 2) to both the raw and phase-flipped channels. This preprocessing pipeline results in 8-channel images going into the network.

For orientation representation, particle rotations are encoded as rotation matrices and mapped to their nearest neighbours in a predefined SO(3) grid. Formally, given a grid **G** of rotation matrices over SO(3) and a point symmetry group Ω, the mapping is defined as:

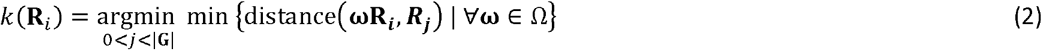

During training, the orientation labels_**L**_*i*_ ∈ ℝ_^|**G**|^ are encoded as sparse vectors containing |Ω| nonzero values, with the remaining elements set to zero.

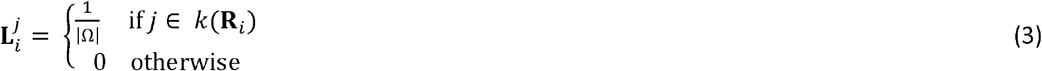

To ensure that the less reliable labels are not used during training, by default, we also exclude particles with the RELION metadata value rlnMaxValueProbDistribution below 0.01 from the training set.

### 4.8. Deep learning model

We employ an upgraded version of our previously published deep learning model^28^ (see Supplementary Table 10 for a performance comparison). The model builds upon the Image2Sphere architecture^42^. It uses first an encoder network to compute N-dimensional 2D features for the input image. These 2D features are then orthographically projected onto a 3D hemisphere. The projected features are interpreted as spherical harmonics and subsequently processed with a group equivariant convolution layer with support in S^2^, followed by a group equivariant convolution with SO(3) support. Finally, the output of the SO(3) convolution, which takes the form of Wigner D-matrix coefficients, is projected into a discretized representation of SO(3) with 294,912 rotation matrices (corresponding to a discretization of ∼3.7°, HEALPix order 4^43^).

In our present implementation, we replace the original ResNet encoder^44^ with a U-Net^45^ with 5 encoder blocks and 4 decoder blocks (see Supplementary Table 3 for a comparison). This allows for larger spatial dimensions in the 2D feature map representing the image, which should lead to better expression upon projection onto the hemisphere.

We also refined the method for estimating the output rotation matrix predR_*i*_. The original implementation directly selected the rotation matrix with the highest probability from a discretized SO(3) grid:

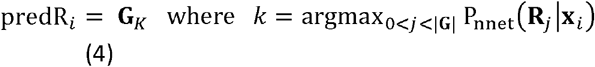

where, G = {R_*j*_} is the set of rotation matrices in the grid that sample SO(3), and P_nnet_ (R_*j*_|x_*i*_) is the model-estimated probability that R_*j*_ represents the pose of particle ***x***_*i*_. In our current approach, we compute the rotation matrix as a weighted average over neighbouring grid points. Specifically, the predicted rotation matrix predR_*i*_ is calculated as:

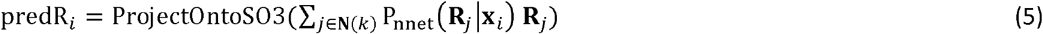

where **N**(*k*) represents the neighbour grid indices of the *k*-th index, ProjectOntoSO3(**M**) = **UV**^T^, and **U** and **V** are obtained via Singular Value Decomposition **M = U Σ V^T^**

We further improved the method by symmetrizing predictions according to the protein point symmetry:

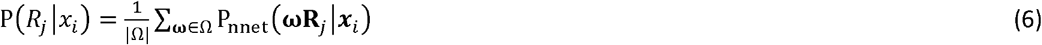

where Ω is the set of rotation matrices given a point symmetry group. As in our previous work, we employ a weighted cross-entropy loss function:

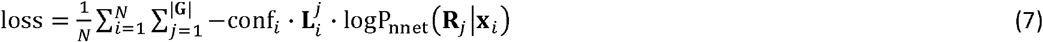

where conf_*i*_ represents the pose reliability estimate given by rlnMaxValueProbDistribution RELION score and 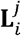 is the label value for the *i*-th particle and *j*-th rotation matrix (see Methods Section 4.7). In our current implementation, we have enhanced the loss function by incorporating label smoothing (*ϵ* = 0.05)^46^ to mitigate overfitting and account for potential inaccuracies in the data labels. Additionally, we implement gradient clipping (default value of 5, value-based) to ensure stable training dynamics.

For a comprehensive overview of all model hyperparameters, please refer to Supplementary Note 2.

### 4.9. Local refinement implementation

A custom PyTorch-based^47^ implementation of the template matching algorithm^10^ has been used in this work as a proof of concept to illustrate the capabilities of our neural network. The algorithm begins by generating, for each particle **x**_*i*_, a grid of orientations (**G**) around the Euler angles predicted by the neural network 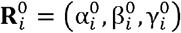.

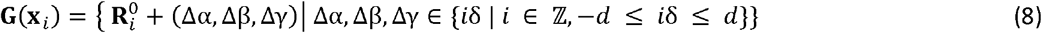

where 2*d* is the grid width (12° by default), and *δ* is the angular step size (2° by default).

Then, for each orientation in the global pool of orientations, the reference volume is projected using the `torch-fourier-slice` Python package^48^. Next, for each particle associated to the orientation, the cross-correlation (CC) between the particle and the CTFs modulated projection is computed.

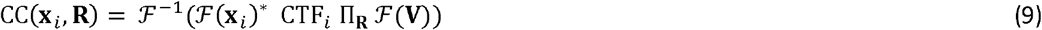

where *V* represents the reference volume, Π_*R*_ is the projector operator at orientation *R*, CTF_*i*_ is the e contrast transfer function of the *i*-th particle and ℱ and ℱ^-1^ represents the Fourier transform and inverse Fourier transform respectively. For efficiency, the Fourier transform of the volume and the particles, as well as CTFs of the particles are precomputed.

After that, the orientation of a particle is assigned to the orientation that yielded the highest cross-correlation peak:

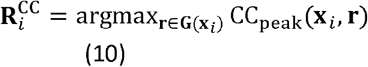

where

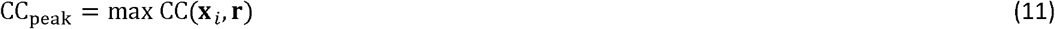

and the shifts of the particle are estimated as the location of the highest cross-correlation peak

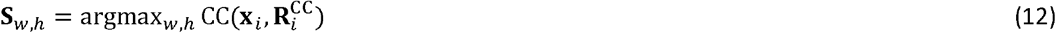

where *w,h* represent the pixel coordinates that are at a distance smaller than 20% of the particle radius from the centre of the image. Finally, a weight for the particle to be used at reconstruction is computed by combining the score given by the neural network and an estimation of the significance of the orientation in a similar fashion as Sorzano et al.^49^:

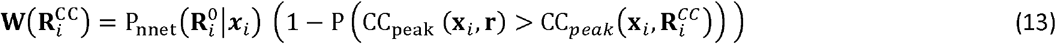

where 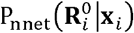 is the probability given by the neural network to the pose 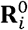 and P(CC_*peak*_(**x**_*i*_,**r**)) is the probability of a cross-correlation peak value for the *i*-th particle at a given pose. For computational reasons, P(CC_*peak*_(**x**_*i*_,**r**)) is assumed to follow a normal distribution whose parameters are estimated as

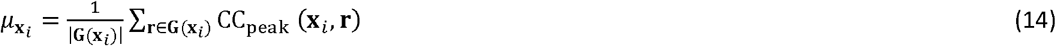

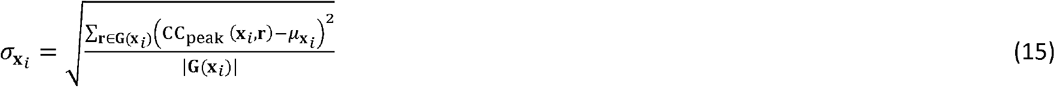

where |G(**x**_*i*_)| is the number of orientations in the grid of orientations for the particle **x**_*i*_

A slower but more accurate two-stage coarse-to-fine search strategy is also implemented. The search consists of two consecutive projection-matching stages using the same cross-correlation metric as in the single-stage approach.

In the first stage, a coarse angular search is performed over a range of ±6° with a step size of 2°, and the top-*K* candidate orientations (default *K* = 5) are retained. In the second stage, a local refinement is carried out independently around each of these *K* candidates, using a narrower search range of ±2.1° with a step size of 0.7°.

After identifying the best discrete orientation on the refinement grid, sub-grid accuracy is achieved via parabolic interpolation along each Euler angle axis. For each axis (rot, tilt, psi), a 1D quadratic function is fitted to the correlation values at the peak and its two neighbouring grid points. The offset from the peak is estimated as:

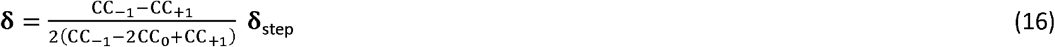

where **CC**_0_ is the correlation at the peak orientation, **CC**_-1_ and CC_+1_ are the correlations at the neighboring grid points along the axis, and **δ**_step_ is the angular step size. The refined Euler angles are then given by:

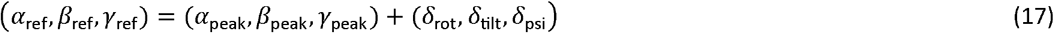

In addition, in-plane shifts are refined beyond integer-pixel accuracy using the same parabolic interpolation scheme applied independently along each image axis. For a correlation peak located at integer coordinates_(*i*_peak_,*j*_peak_)_, the sub-pixel offsets are computed as:

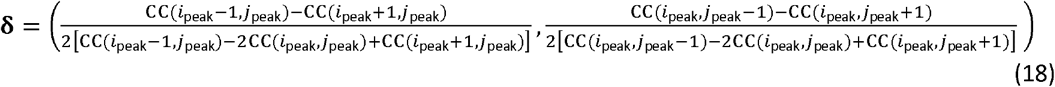

The final sub-pixel shift estimate is given by:

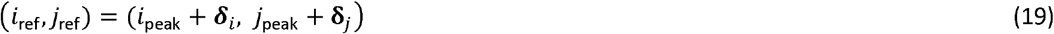

For a sensitivity analysis of the main parameters of this algorithm, see Supplementary Note 4 and Supplementary Table 5.

### 4.10. Particle pruning

After the neural network has predicted a probability distribution for the orientations of a particle *xi*, we compute its direction-normalized pruning score as a robust z-score based on the median absolute deviation (MAD) of the distribution of predicted scores at a given orientation obtained from the validation set at training time. Particles with a robust z-score below a given threshold are regarded as “bad” particles and ignored in subsequent steps.

The median and MAD calculations required to compute our directional robust z-scores can be regarded as part of the training process and are computed as follows. First, the particles in the validation set are assigned to one of the *K* cones in which we discretize the projection sphere (first and second Euler angles according to RELION’s convention). By default, we use cones of 15° (HEALPix order of 2) to ensure a sufficient number of particles in each cone. Then, the cone median and MAD are computed as

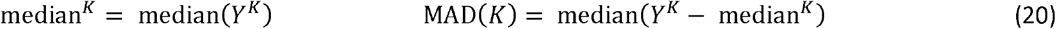

where

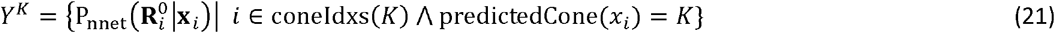

Here, coneIdxs (*K*) is the set of particle indices that correspond to the cone *K* according to the ground truth pose, and predictedCone(**x**_*i*_) is the cone index of the highest score orientation of the and estimated from the validation dataset are then used to compute the *i*-th particle.

The median^*K*^ and MAD (*K*) estimated from the validation dataset are then used to compute per-cone robust z-score of new particles following the next equation:

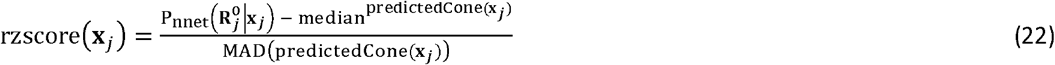

In this study, for robust-z score calculations, we define the particle score 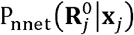 as the sum of the top-10 highest probabilities predicted by the neural network. This approach makes the score more robust in cases where the neural network is uncertain about particle orientation, as the top-1 probability might not capture enough of the learnt signal.

There are several approaches to estimate a per-cone robust z-score threshold. For instance, it could be computed based on the expected fraction of “bad” picks given historical data. Here we estimate it by comparing the distribution of per-cone robust z-scores of the validation dataset against a dataset of “bad” particles (particles that were removed after 2D classification). See Supplementary Note 3 and Supplementary Fig. 2-4 for details about automatic thresholds.

## 5. Data Availability

The cryo-EM maps generated in this study have been deposited in the Electron Microscopy Data Bank (EMDB) under accession codes EMD-55146 (β-galactosidase–B1), EMD-55241 (β-galactosidase– B2), EMD-55304 (PKM2–P1), EMD-55305 (PKM2–P2), EMD-55535 (GDH–G1), EMD-55549 (GDH–G2), EMD-55532 (GDH–G3), and EMD-58344 (TRPML1-T1). The atomic coordinates generated in this study have been deposited in the PDB under accession codes 9T4W (G1), 9T4X (G2), and 9T4U (G3). The raw particle images generated in this study are available in the Electron Microscopy Public Image Archive (EMPIAR) under accession numbers EMPIAR-13104 (G1), EMPIAR-13105 (G2), and EMPIAR-13106 (G3). Source data are provided with this paper.

## 6. Code availability

CryoPARES is available at https://github.com/rsanchezgarc/cryoPARES with DOI https://doi.org/10.5281/zenodo.21725717.

## 8. Acknowledgments

This work was part of a Sustaining Innovation Research Programme at Astex Pharmaceuticals. R.S.-G. was a Sustaining Innovation Postdoctoral Research Associate at Astex Pharmaceuticals and thanks Astex Pharmaceuticals for funding. We acknowledge the Cambridge Pharmaceutical Cryo-EM Consortium and the Astbury BioStructure Laboratory for access to their cryo-EM facilities.

## 9. Author contributions

R.S.G. co-designed, implemented, and tested the method, performed image processing and wrote the first draft of the manuscript. A.A., A.B., J.R., and P.A.W. prepared the protein samples and collected electron microscopy data. C.P. co-designed the method. M.S. prepared the training and testing data, analysed the results, and co-supervised the project. C.D. co-supervised the project and secured funding. All authors contributed to the writing of the final version of the manuscript.

