## Supplementary Information for "Supervised Deep Learning for Efficient Cryo-EM Image Alignment in Drug Discovery with cryoPARES"

### CryoPARES supplementary information

#### 1. Supplementary Figures

##### Training curves for simulated and trained-from-scratch experiments

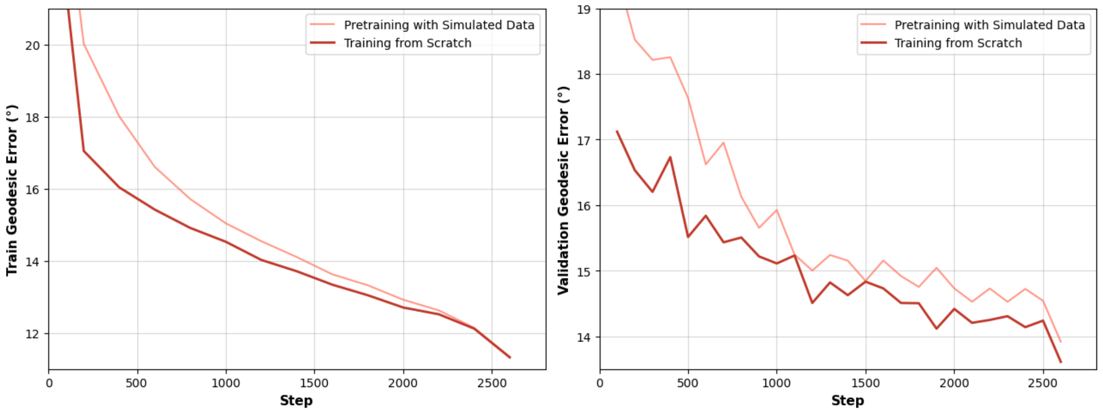

Supplementary Fig. 1. Training curves recording the mean geodesic error for the training set (left) and validation set (right) for the PKM2 apo dataset using the simulation pretraining (red), and training from scratch using experimental particles only (orange). Particle simulation was performed using the `relion_project` command with the following options `--simulate --adjust_simulation_SNR 2.0 --ctf`.

#### Calibration of the thresholds for the direction-normalized pruning scores

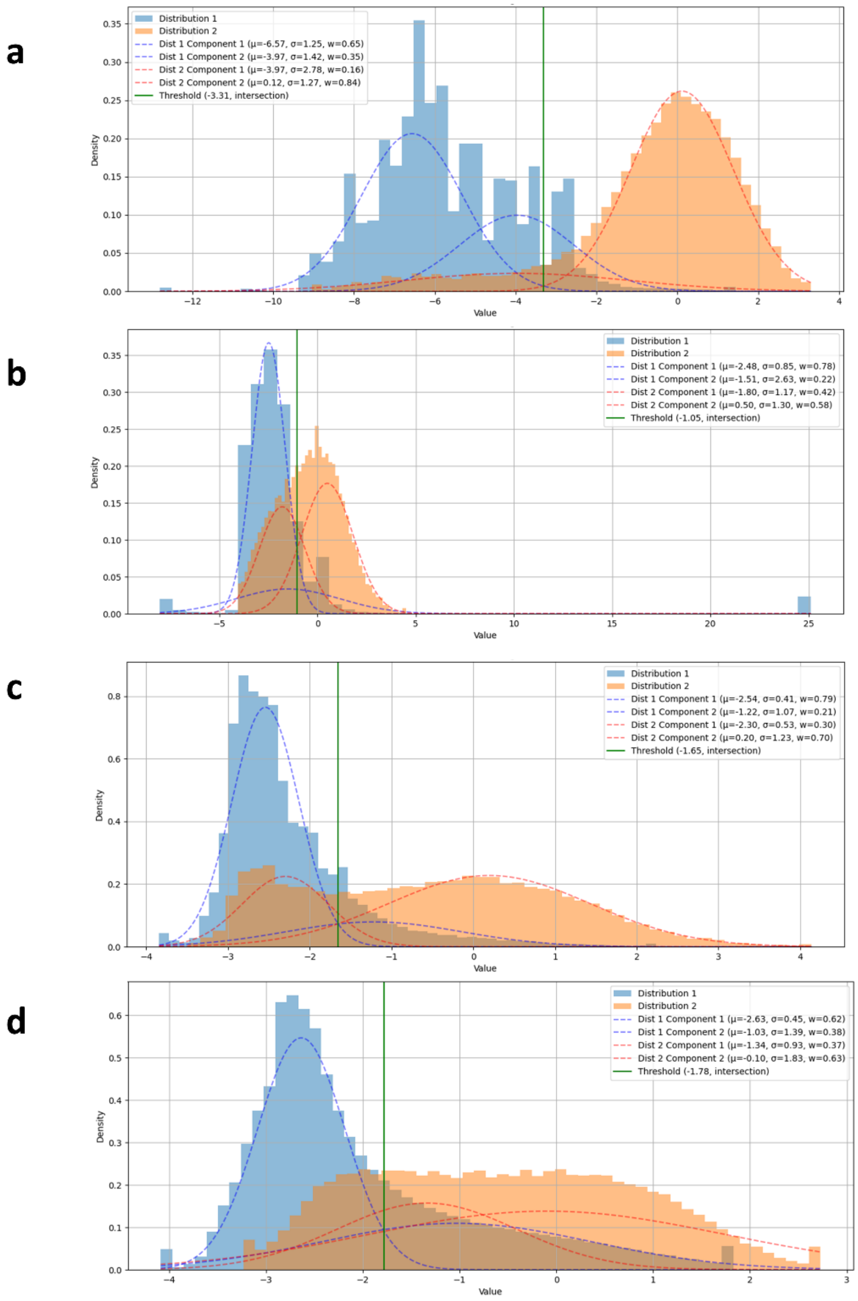

Supplementary Fig. 2. Distribution of direction-normalized pruning scores for the validation set (orange) of the target proteins used for training (a, BGAL; b, PKM2; c, GDH; d, TRPML1), compared to the distribution of pruning scores for "bad" particles (blue), defined as particles that were rejected after a 2D classification run. Hashed lines represent the components of the Gaussian Mixture Model (GMM) with two components fitted to the distributions. The green vertical lines mark the selected threshold based on the intersection of the GMM component with smaller mean from the distribution of "bad" particles and the component with larger mean from the distribution of "good" particles. μ, σ, and w refer to the mean, standard deviation and weight of each of the components of the GMM.

**Confidence of the pose prediction for different viewing cones**

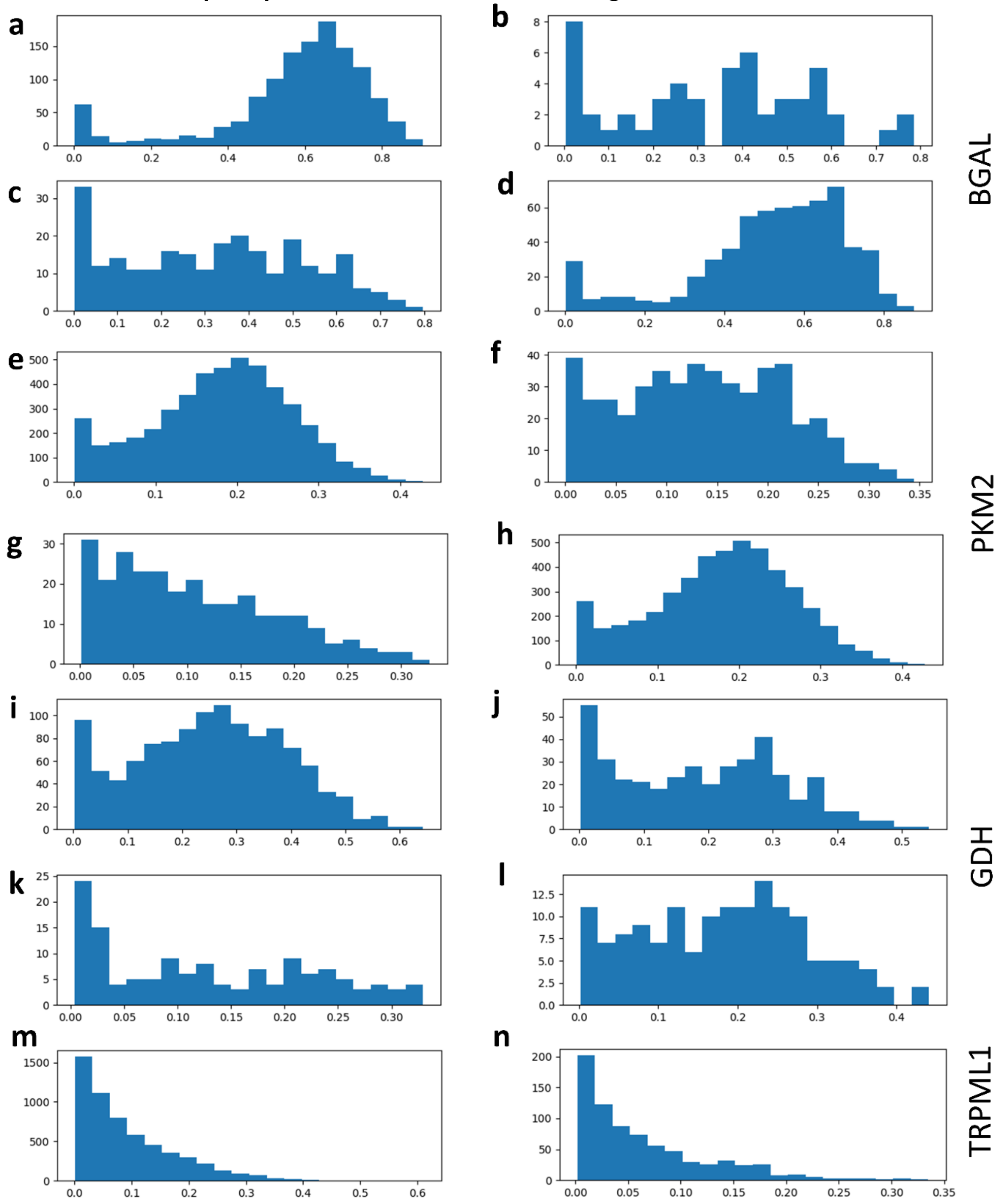

Supplementary Fig. 3. Distribution of neural network top-1 confidence predictions (range 0 to 1) for particles at different projection cones for BGAL (a-d), PKM2 (e-h), GDH (i-l) and TRMPL1 (m-n). The cones encompass particles up to 7.5° from the centre. The cones shown were manually selected to illustrate the variation in distributions between different cones, highlighting the importance of a directional normalization.

#### Pruning score distributions for good and bad particles

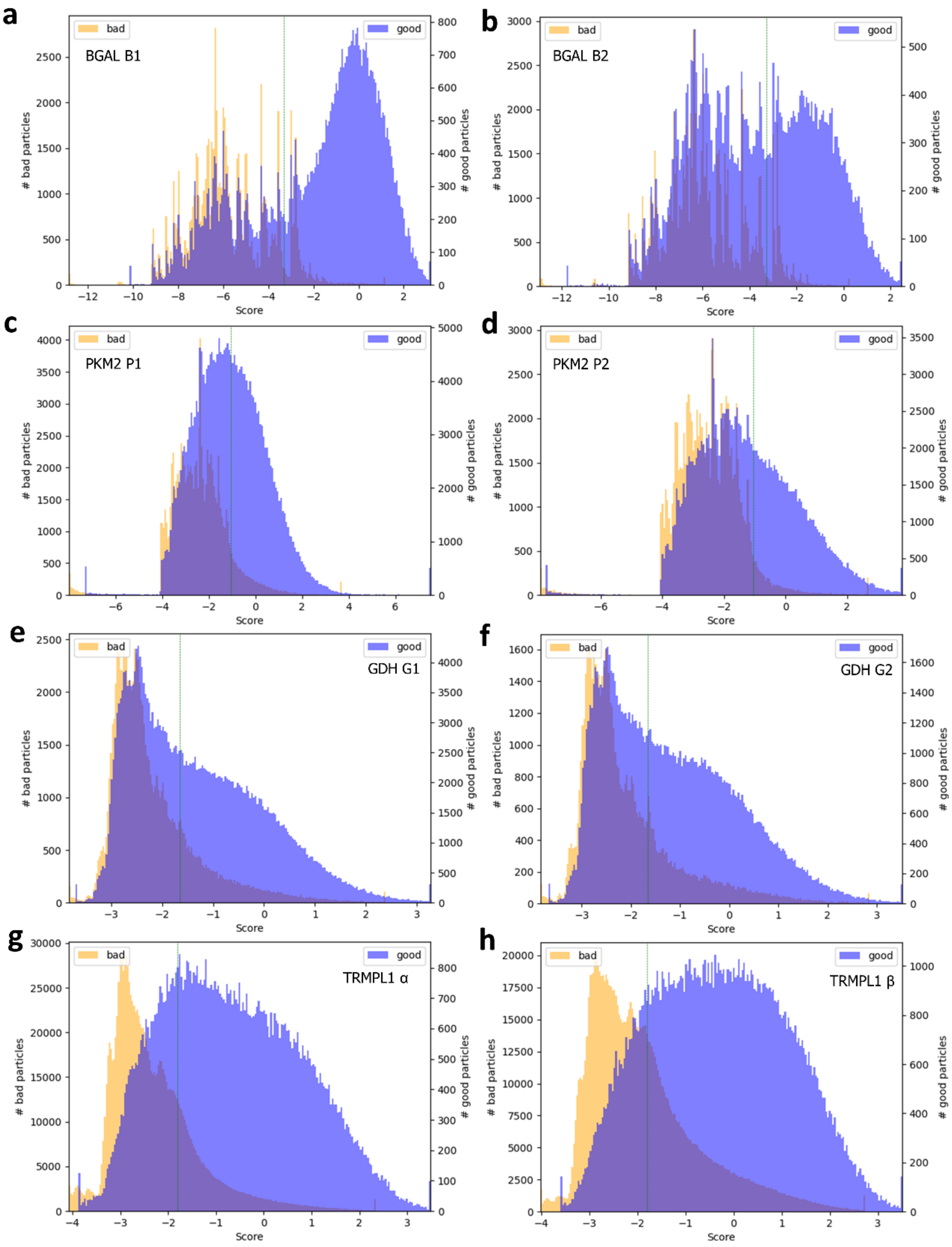

Supplementary Fig. 4. Histograms showing the distribution of direction-normalized pruning scores for particles that were discarded after 2D classification (bad, orange), and particles that survived the 2D classification step (good, blue). Note the different scales for number of “good” and “bad” particles. (a) BGAL bound to Compound B1, (b) and Compound B2, (c) PKM2 bound Compound P1, (d) and Compound P2, (e) GDH bound to Compound G1, (f) and Compound G2, (g) membrane protein bound to ligand TRPML1 α and β (h). The vertical green line corresponds to the threshold automatically estimated for each target (see main text Methods Section 4.8).

**2D class-averages from cryoPARES pruned particles**

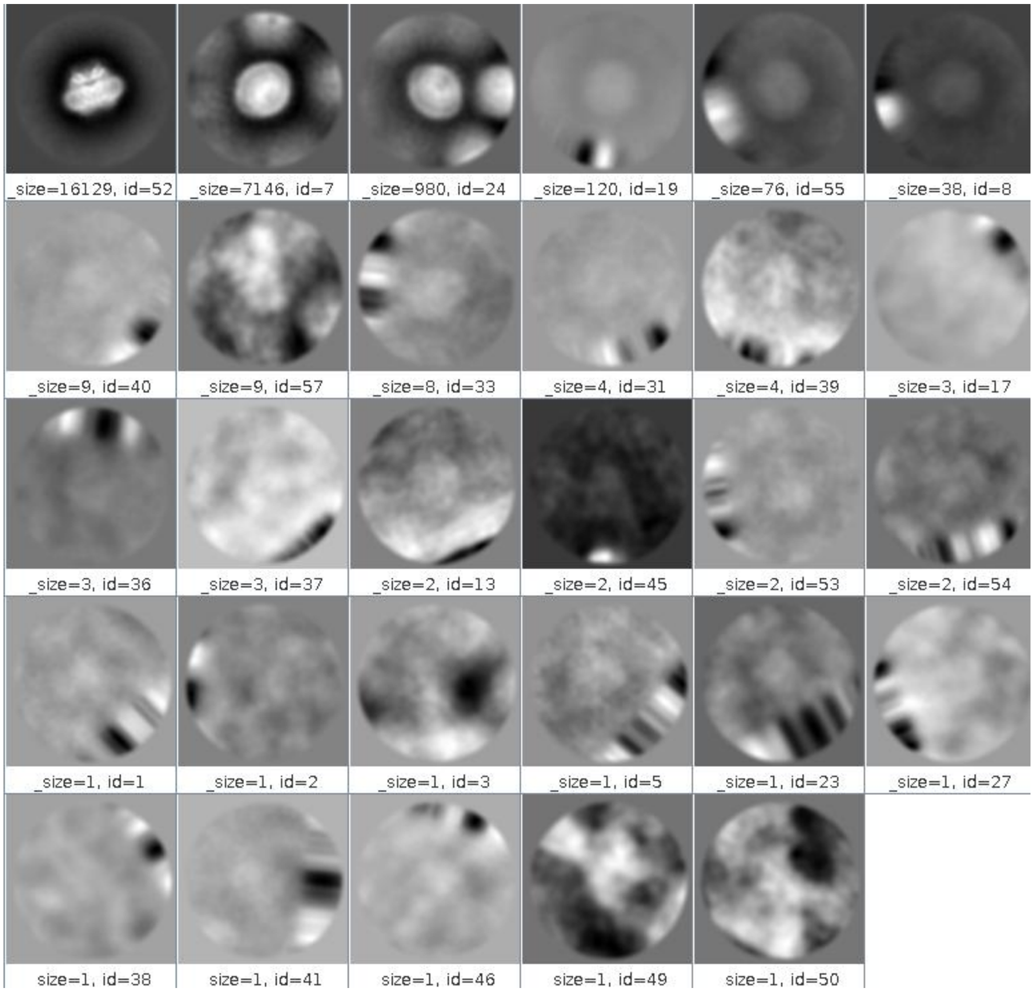

Supplementary Fig. 5. 2D class averages from a RELION 2D classification job run on the particles rejected by cryoPARES pruning in the TRPML1 T1 dataset. The majority of classes show featureless or heterogeneous density consistent with junk particles. One class displays recognizable TRPML1 structural features, indicating that a fraction of the rejected particles are morphologically intact protein particles whose pose estimates did not meet the cryoPARES confidence threshold. Despite being structurally recognizable, the predicted poses of these particles exhibit a median angular error of 26.7° relative to RELION reference poses, compared to 3.7° for the cryoPARES-retained particles, confirming that their rejection is justified on the basis of pose reliability.

**FSC curves for different model bias experiments**

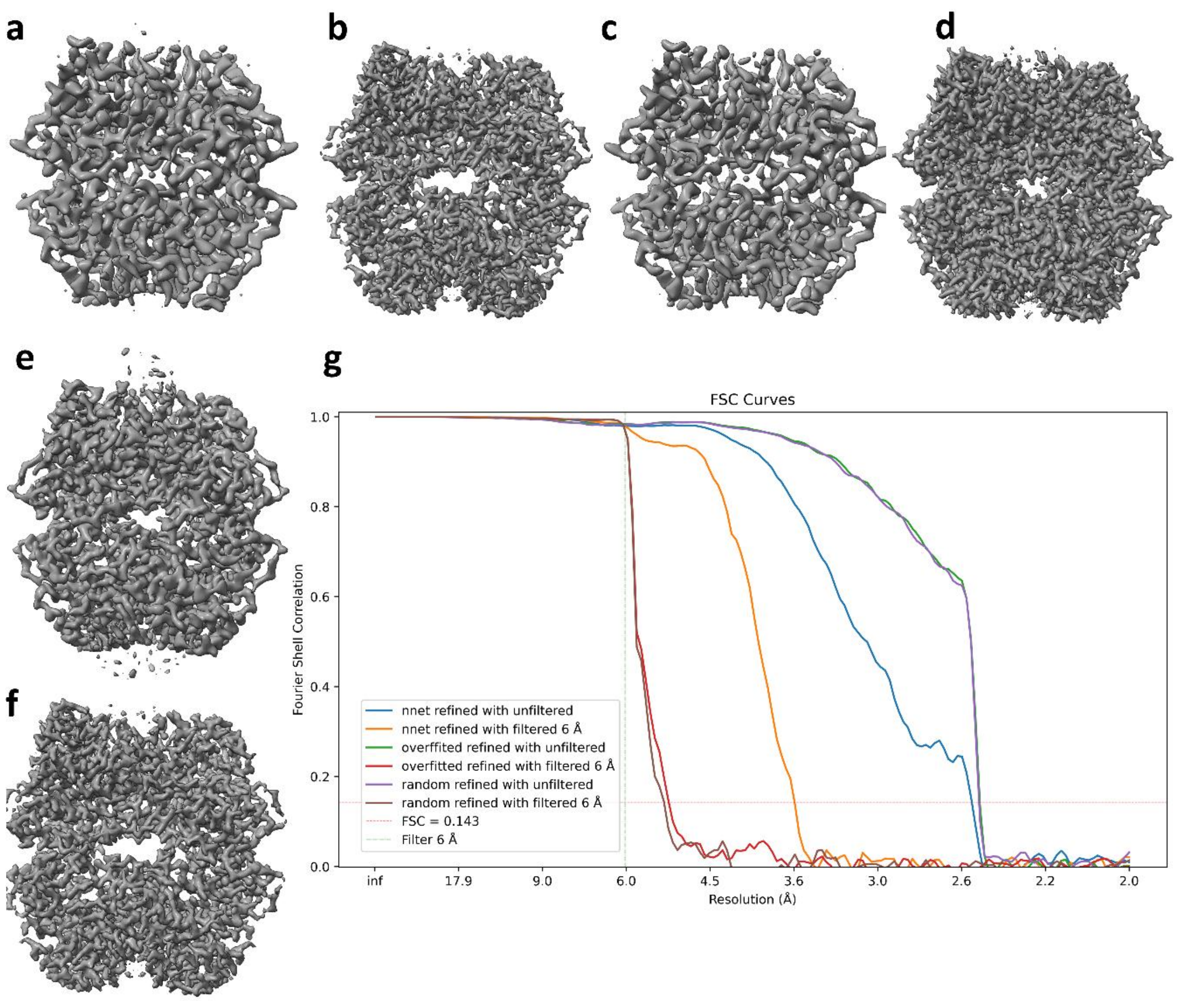

Supplementary Fig. 6. Reconstructed PKM2 Compound P1 under different model bias situations. a) Map obtained after applying a local refinement step to a set of particles with randomised orientations when the template reference is low-pass filtered at 6 Å, and b) when an unfiltered template is used instead. c) Map obtained when a local refinement step is applied to the already overfitted angular assignments of (b) if the reference map is filtered at 6 Å or d) left unfiltered. e) Map obtained after applying a local refinement step to a set of particles with cryoPARES orientations when the template reference is low-pass filtered at 6 Å, and f) when an unfiltered template is used instead. g) FSC curves for the experiments a-f. The vertical hashed, green line shows the low-pass cut-off of the filtered references.

**Map anisotropy estimation for the ADP-bound GDH reconstructions**

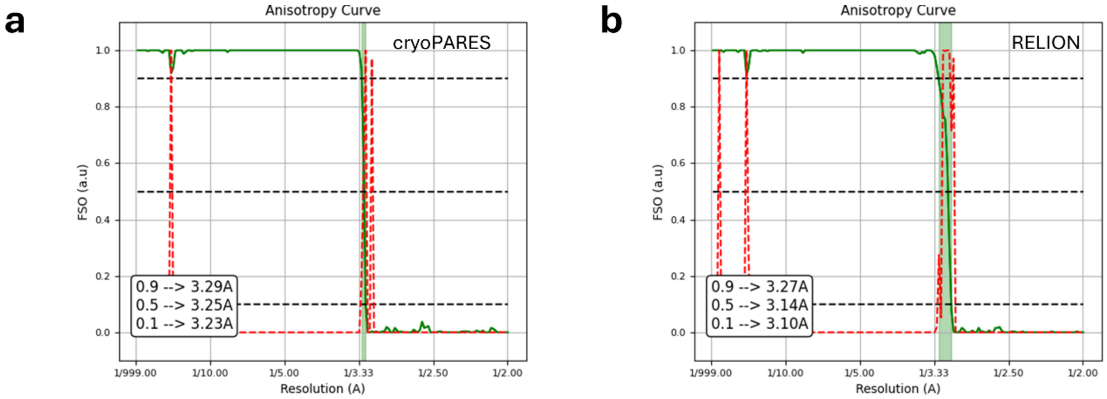

Supplementary Fig. 7. Map-to-model FSO curves for GDH G3 reconstructions obtained with cryoPARES (a) and RELION (b; reconstruction performed using a 15 Å low-pass filtered reference). The cryoPARES map exhibits a narrower FSO percentile range (shadowed region), indicating reduced anisotropy compared to RELION. The atomic model used for FSO calculation was derived from an independent ADP-bound GDH dataset that did not exhibit preferred orientation.

**Alternative visualization of the ligands showed in Main Text Figures 2-5**

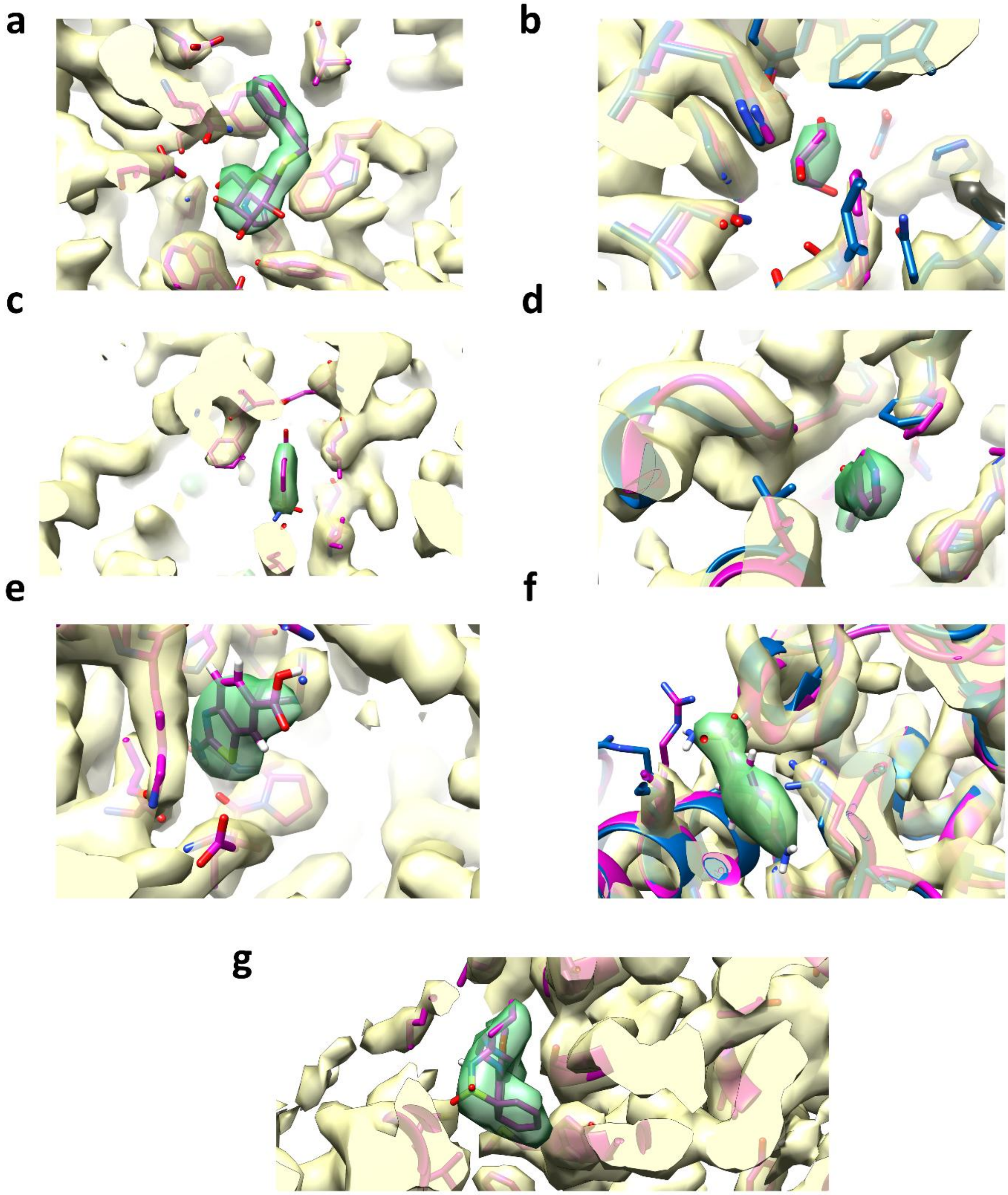

Supplementary Figure 8. Alternative visualization of the ligands showed in Main Text Figures 2-5 using the same contour level. BGAL bound to ligand B1 (a) and B2 (b), PKM2 bound to ligand P1 and P2, GDH bound to ligand G1 and G2, and TRPML1 bound to ligand T1. The density closer than 1.8 Å to the ligand is coloured in green.

**Local Resolution of the maps reconstructed from cryoPARES-estimated poses**

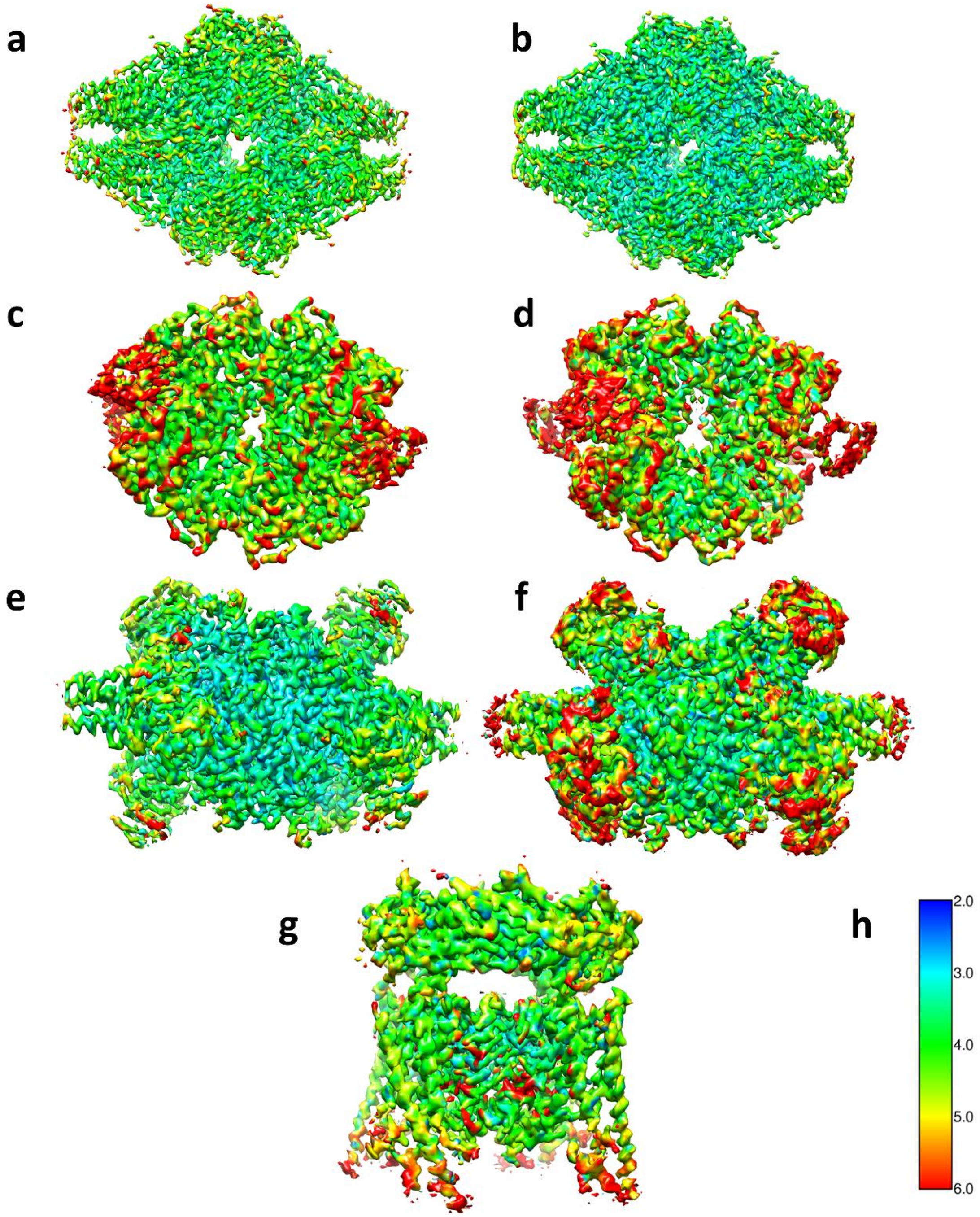

Supplementary Fig. 9. Maps reconstructed from cryoPARES-estimated orientations coloured by local resolution estimated with Monores[1]: BGAL bound to Compound B1 (a) and Compound B2 (b), PKM2 bound to Compound P1 (c) and Compound P2 (d), GDH bound to Compound G1 (e) and Compound G2 (f), TRPML1 bound to Compound T1 (g). The colour bar showing the resolution in Å is displayed in f.

**GDH datasets image processing workflow**

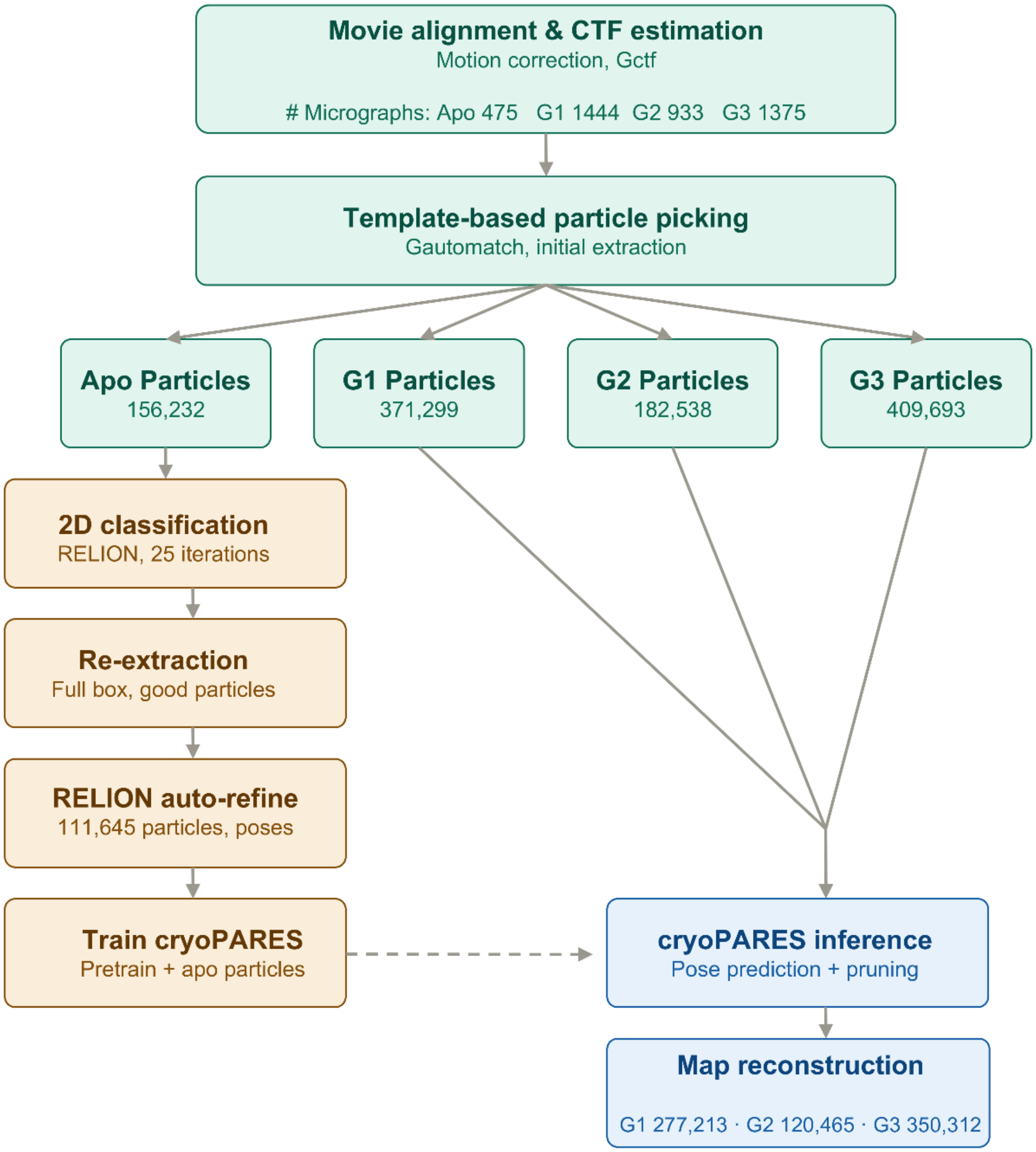

Supplementary Fig. 10. Image processing workflow for the GDH datasets. The number of micrographs for each sample is included in the topmost box. The number of particles obtained after each step is included in their corresponding boxes.

**Representative 2D classes for the GDH datasets**

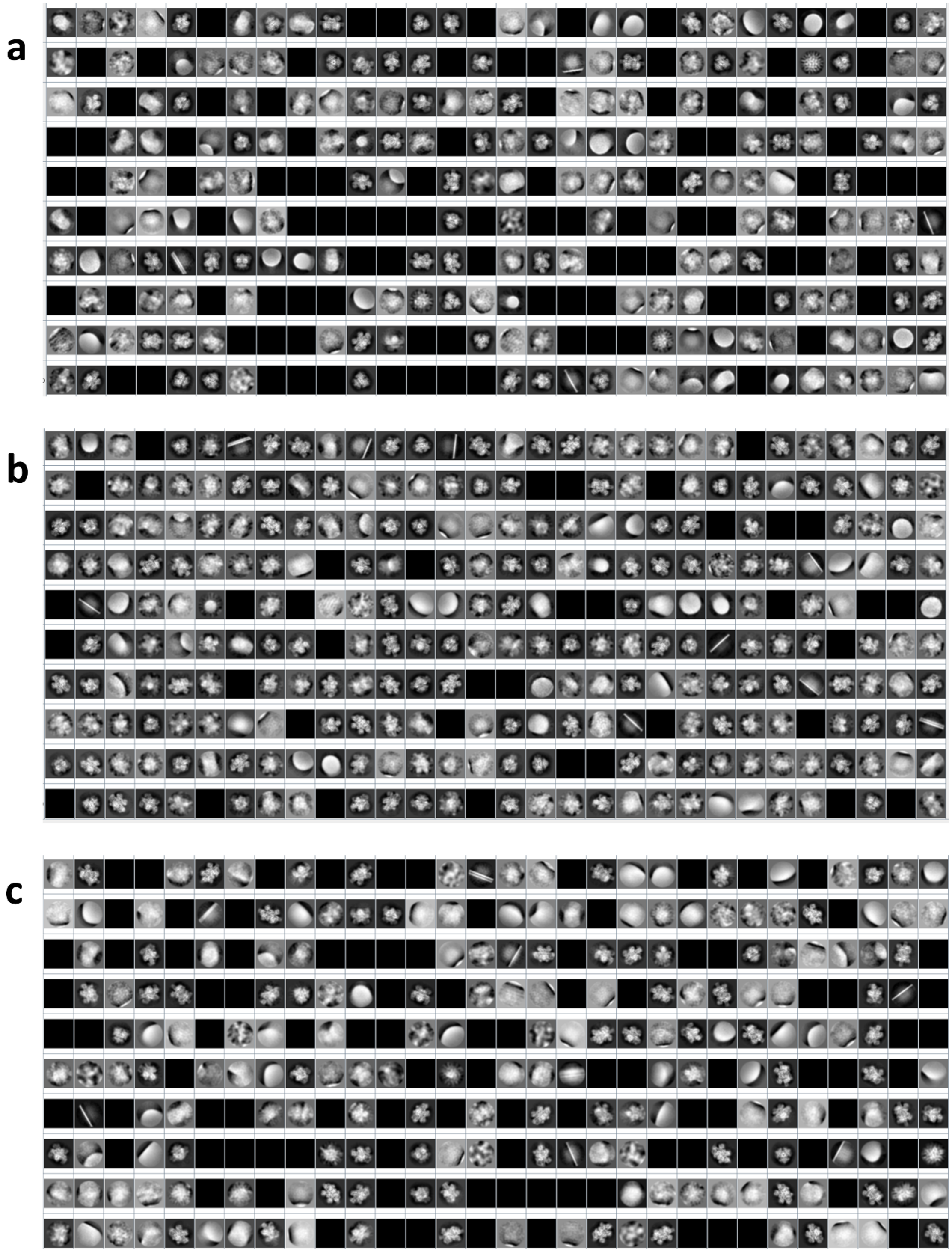

Supplementary Fig. 11. RELION 2D class averages for the datasets GDH apo (a), GDH G1 (b), and G2 (c).

**Representative micrographs and angular distributions for the GDH datasets**

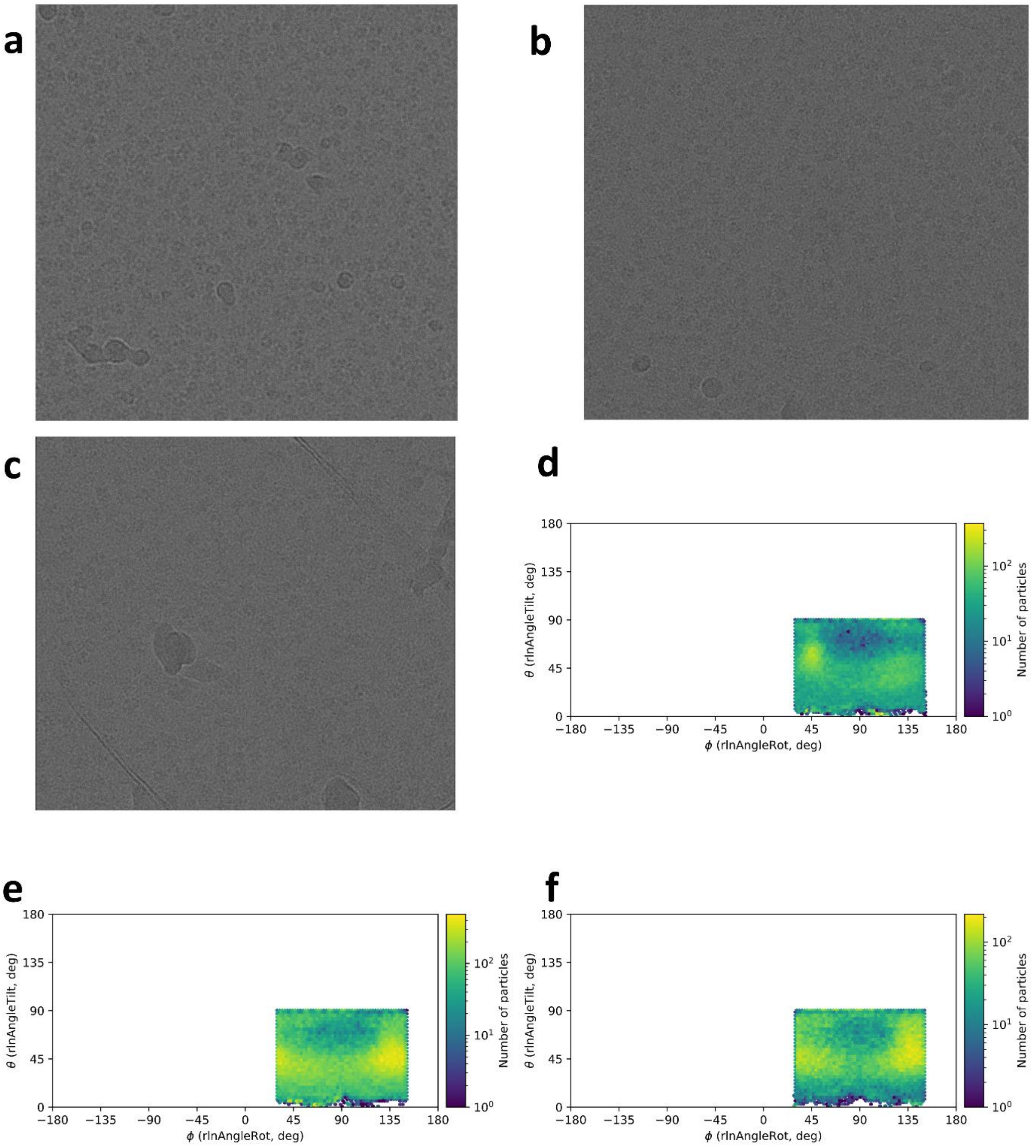

Supplementary Fig. 12. Representative micrographs for the datasets GDH apo (a), GDH G1 (b), and G2 (c) and angular distributions for the particle poses of the same datasets (GDH apo, d; GDH G1, e; and G2, f). Note that the D3 symmetry of the sample is explicitly included in the panels d-f.

#### 2. Supplementary Tables

Supplementary Table 1. Cryo-EM data collection, refinement and validation statistics for the targets studied. Atomic models for BGAL, PKM2, and TRPML1 were directly downloaded from the PDB and rigid-body fitted into the densities. GDH models were built from the maps.

| | BGAL apo (EMDB-55522) | BGAL B1 (EMDB-55146) | BGAL B2 (EMDB-55241) | PKM2 apo (EMDB-55516) | PKM2 P1 (EMDB-55304) | PKM2 P2 (EMDB-55305) | GDH apo (EMDB-55559) | GDH G1 (EMD-55535) (PDB 9T4W) | GDH G2 (EMD-55549) (PDB 9T4X) | GDH G3 (EMD-55532) (PDB 9T4U) | TRPML1 T1 (EMD -58344) (PDB 9HLB) |
|---|---|---|---|---|---|---|---|---|---|---|---|
| **Data collection and processing** | | | | | | | | | | | |
| Magnification | 120k x | 120k x | 120k x | 120k x | 190k x | 120k x | 120k x | 120k x | 120k x | 120k x | 120k x |
| Voltage (kV) | 300 | 300 | 300 | 200 | 200 | 300 | 300 | 300 | 300 | 300 | 300 |
| Electron exposure (e–/Å$^2$) | 69.015 | 65.53 | 66.83 | 66.4275 | 54.85 | 65.4 | 70.58 | 62.51 | 62.51 | 70.05 | 39.08 |
| Defocus range (μm) | 1.0-3.0 | 0.7-2.5 | 0.7-2.1 | 0.8-2.2 | 0.7-2.1 | 0.8-2.2 | 0.8-2.2 | 0.8-2.2 | 0.8-2.2 | 0.8-2.2 | 0.8-2.0 |
| Pixel size (Å) | 0.674 | 0.674 | 0.674 | 0.674 | 0.752 | 0.674 | 0.674 | 0.6788 | 0.6788 | 0.666 | 0.6431 |
| Symmetry imposed | D2 | D2 | D2 | D2 | D2 | D2 | D3 | D3 | D3 | D3 | C4 |
| Number of movies collected | 1552 | 562 | 517 | 569 | 707 | 590 | 475 | 1444 | 933 | 1375 | 6381 |
| Initial particle images (no.) | 369,690 | 134,171 | 124,687 | 212,244 | 353,678 | 267,921 | 156,232 | 371,299 | 182,538 | 409,693 | 2,248,589 |
| Final particle images (no.) | 177,824 | 56,764 | 42,125 | 125,324 | 254,070 | 127,035 | 111,645 | 277,213 | 120,465 | 350,312 | 398,353 |
| Map resolution (Å) at 0.143 FSC threshold | 2.42 | 2.8 | 3.2 | 2.6 | 3.5 | 3.3 | 2.58 | 2.9 | 3.0 | 3.6 | 2.7 |

| | | | | | | | | | | | |
|---|---|---|---|---|---|---|---|---|---|---|---|
| Map resolution range (Å) | - | - | - | - | - | - | - | - | - | - | |
| **Refinement** | | | | | | | | | | | |
| Initial model used (PDB code) | - | 6TTE | 6TSK | - | 6TTF | 6TTQ | - | 3JCZ | 3JCZ | 3JCZ | 9HLB |
| Model resolution (Å)<br>FSC threshold | -<br>- | -<br>- | -<br>- | -<br>- | -<br>- | -<br>- | -<br>- | 2.87<br>0.5 | 3.1<br>0.5 | 3.63<br>0.5 | -<br>- |
| Model resolution range (Å) | - | - | - | - | - | - | - | - | - | - | - |
| Map sharpening *B* factor (Å$^2$) | | | | | | | | -103.38 | -100.79 | -125.74 | - |
| Model composition<br>Non-hydrogen atoms<br>Protein residues<br>Ligands | -<br>-<br>- | -<br>-<br>- | -<br>-<br>- | -<br>-<br>- | -<br>-<br>- | -<br>-<br>- | -<br>-<br>- | 23370<br>2958<br>18 | 23346<br>2952<br>18 | 23304<br>2958<br>6 | -<br>-<br>- |
| *B* factors (Å$^2$)<br>Protein<br>Ligand | -<br>- | -<br>- | -<br>- | -<br>- | -<br>- | -<br>- | -<br>- | 72.92<br>50.89 | 82.52<br>71.98 | 92.00<br>68.91 | -<br>- |
| R.m.s. deviations<br>Bond lengths (Å)<br>Bond angles (°) | -<br>- | -<br>- | -<br>- | -<br>- | -<br>- | -<br>- | -<br>- | 0.0072<br>1.02 | 0.0067<br>0.96 | 0.0054<br>0.95 | -<br>- |
| Validation<br>MolProbity score<br>Clashscore<br>Poor rotamers (%) | -<br>-<br>- | -<br>-<br>- | -<br>-<br>- | -<br>-<br>- | -<br>-<br>- | -<br>-<br>- | -<br>-<br>- | 2.2<br>6.1<br>3.51 | 1.4<br>4.32<br>0.08 | 1.64<br>5.42<br>0.2 | -<br>-<br>- |
| Ramachandran plot<br>Favored (%) | -<br>- | -<br>- | -<br>- | -<br>- | -<br>- | -<br>- | -<br>- | 67.55<br>28.93 | 81.59<br>18.33 | 75.67<br>24.13 | -<br>- |

| | | | | | | | | | | | |
|---|---|---|---|---|---|---|---|---|---|---|---|
| Allowed (%)<br>Disallowed (%) | - | - | - | - | - | - | - | 3.51 | 0.08 | 0.2 | - |

Supplementary Table 2. Compounds employed in this work.

| Compound ID | Target | EMPIAR ID | Name | 2D structure |
|---|---|---|---|---|
| B1 | BGAL | 10644 | PETG | 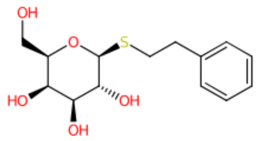 |
| B2 | BGAL | 10646 | L-ribose (pyranose form) | 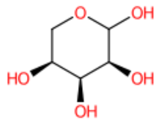 |
| P1 | PKM2 | 10648 | 5-hydroxynaphthalene-1-sulfonamide | 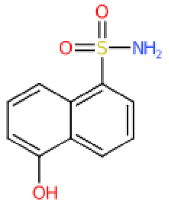 |
| P2 | PKM2 | 10649 | 1-propan-2-yl-3-pyridin-4-yl-urea | 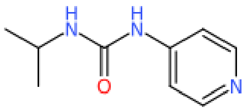 |
| G1 | GDH | 13104 | 2-amino-1,3-benzothiazole-6-carboxylic acid | 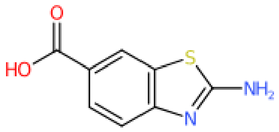 |
| G2 | GDH | 13105 | 2-amino-1,3-benzothiazole-6-sulfonamide | 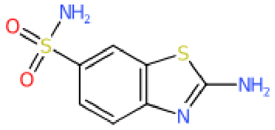 |
| G3 | GDH | 13106 | ADP | 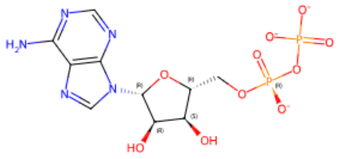 |

| | | | | |
|---|---|---|---|---|
| T1 | TRPML1 | NA | N-[2-[4-(2-methoxyphenyl)piperazin-1-yl]phenyl]benzenesulfonamide | 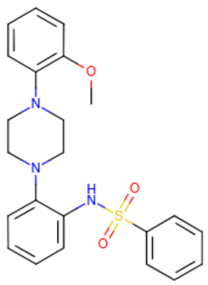 |

Supplementary Table 3. ResNet vs UNet validation performance comparison. The metrics were computed on the validation set used during training.

| Dataset | Model | Validation mean geodesic error (°) | Validation median geodesic error (°) |
|---|---|---|---|
| BGAL | ResNet101 | 15.7 | 6.9 |
| | UNet | 13.7 | 5.5 |
| PKM2 | ResNet101 | 8.9 | 3.8 |
| | UNet | 7.8 | 3.1 |
| GDH | ResNet101 | 22.3 | 7.0 |
| | UNet | 20.5 | 5.6 |

Supplementary Table 4. Comparison of particle orientation angles across different methods. We report the median geodesic distance (medG) and the percentage of particles with orientations within 5° or 10° between: (1) RELION auto-refine estimates versus raw neural network predictions, (2) RELION versus neural network predictions followed by local refinement (±6° search range, 2° steps), and (3) two independent RELION auto-refine jobs.

| | Nnet vs RELION | | | Nnet + local refinement vs RELION | | | RELION vs RELION | | |
|---|---|---|---|---|---|---|---|---|---|
| | medG | %< 5° | %< 10° | medG | %< 5° | %< 10° | medG | %< 5° | %< 10° |
| BGAL B1 | 4.0° | 61.0% | 75.3% | 1.6° | 75.4% | 77.0% | 0.1° | 86.5% | 88.6% |
| BGAL B2 | 6.0° | 42.4% | 63.1% | 2.2° | 63.3% | 66.8% | 0.2° | 82.3% | 85.0% |
| PKM2 P1 | 7.1° | 30.1% | 65.3% | 3.1° | 65.2% | 74.3% | 1.1° | 86.7% | 90.5% |
| PKM2 P2 | 7.0° | 32.0% | 58.3% | 2.9° | 66.1% | 75.8% | 0.7° | 85.6% | 89.3% |
| GDH G1 | 7.5° | 32.0% | 58.4% | 3.4° | 57.1% | 63.0% | 0.4° | 76.5% | 78.4% |
| GDH G2 | 7.2° | 32.7% | 60.9% | 3.1° | 59.9% | 66.2% | 0.4° | 78.0% | 79.8% |
| TRPML1 T1 | 6.1° | 39.9% | 68.8% | 3.7° | 58.4% | 71.2% | 0.4° | 84.2% | 86.9% |

Supplementary Table 5. Impact of the configuration parameters of the local refinement on the reconstructed maps for the PKM2 P2 dataset.

| Search range | Step | Low-pass filter cutoff | Resolution |
|---|---|---|---|
| ±6° | 2° | 6 Å | 3.28 Å |
| ±6° | 1° | 6 Å | 3.22 Å |
| ±6° | 1° | 4 Å | 3.01 Å |
| ±6° | 1° | 3.5 Å | 2.99 Å |
| ±4° | 0.5° | 3.5 Å | 3.02 Å |
| ±8° | 1° | 4 Å | 2.99 Å |

Supplementary Table 6. CryoPARES throughput measured as particles $min^{-1}$ $GPU^{-1}$ for different samples of different size and different GPUs.

| Target | Symmetry | Image size | Sampling rate | GPU | Nnet inference | Local Refinement 2° | Local Refinement 4° | Local Refinement 6° |
|---|---|---|---|---|---|---|---|---|
| PKM2 P1 | D2 | 334 | 1.0 | A5000 | 10,080.7 | 9,891.5 | 5,895.5 | 2,007.8 |
| PKM2 P1 | D2 | 334 | 1.0 | A100 S-XM4 | 32,690.1 | 15,450.9 | 10,620.1 | 6,000.3 |
| PKM2 P1 | D2 | 222 | 1.5 | A100 S-XM4 | 45,863.8 | 25,045.2 | 18,203.2 | 11,850.9 |
| EMPIAR -10280 | C2 | 182 | 1.5 | A100 S-XM4 | 45,261.3 | 55,481.6 | 39,667.1 | 25,110.5 |
| EMPIAR -10280 | C2 | 182 | 1.5 | A5000 | 28,300.8 | 42,776.4 | 33,559.6 | 12,051.3 |
| EMPIAR -10166 | C1 | 336 | 1.0 | A100 S-XM4 | 31,817.7 | 10,056.2 | 6,351.9 | 4,032.3 |
| EMPIAR -10166 | C1 | 284 | 1.5 | A5000 | 15,325.8 | 17,259.1 | 8,206.1 | 4,787.9 |
| EMPIAR -10166 | C1 | 284 | 1.5 | A100 S-XM4 | 47,186.8 | 18,135.5 | 11,341.2 | 7,363.8 |

Supplementary Table 7. Throughput measured as particles $min^{-1}$ $GPU^{-1}$ for the target PKM2 Compound P1 (254,070 particles, symmetry D2) using RELION auto-refine and cryoSPARC homogeneous refinement.

| Software | Box size | GPU Type | Time (min:s) | #GPUs | Particles $min^{-1}$ $GPU^{-1}$ |
|---|---|---|---|---|---|
| cryoSPARC | 332 | A100 | 08:55 | 1 | 28,493.8 |
| cryoSPARC | 222 | A100 | 08:04 | 1 | 31,496.2 |
| cryoSPARC | 332 | A5000 | 28:04 | 1 | 9,052.3 |
| RELION | 332 | A100 | 99:28 | 4 | 638.5 |
| RELION | 222 | A100 | 89:53 | 4 | 706.6 |
| RELION | 332 | A5000 | 160:00 | 4 | 423.4 |

Supplementary Table 8. CryoPARES training wall time. The training time scales almost linearly with the number of particles.

| Sample | # particles | # Epochs | GPU | Wall time (h) |
|---|---|---|---|---|
| BGAL APO | 177,824 | 55 + 60 | 4 x A5000 | 25.7 |
| BGAL APO | 177,824 | 26 + 29 | 4 x A100 | 18.2 |
| PKM2 APO | 125,324 | 47 + 46 | 4 x A5000 | 13.8 |
| GDH APO | 111,645 | 50 + 52 | 4 x A5000 | 14.5 |
| TRPML1 apo (subset) | 74,626 | 53 + 51 | 4 x A5000 | 8.6 |
| TRPML1 apo | 395,434 | 51 + 49 | 4 x A5000 | 49.1 |
| TRPML1 apo (superset) | 1,298,553 | 57 + 63 | 4 x A5000 | 121.5 |

Note: # Epochs display the epoch number at which the early-stopping callback interrupted the training for each of the half-dataset models. Wall time reports the combined execution time for both half-dataset models. Superset was generated by combining micrographs from different experiments.

Supplementary Table 9. Ω as defined in Lucas et al.[2] and $\Omega^0$ (defined in Supplementary Note 7) values for reconstructions obtained with cryoPARES followed by a local refinement with a template low-pass filtered at 6 Å and an unfiltered template, compared to the values obtained for random orientations followed by a local refinement with an unfiltered reference.

| Sample | cryoPARES Filter 6 Å | | cryoPARES Unfiltered | | Random Unfiltered | |
|---|---|---|---|---|---|---|
| | $\Omega^0$ | $\Omega$ | $\Omega^0$ | $\Omega$ | $\Omega^0$ | $\Omega$ |
| BGAL B1 | 7.1% | 3.6% | 7.1% | 3.4% | 29.8% | 26.7% |
| BGAL B2 | 5.7% | 0.1% | 8.7% | 1.5% | 26.1% | 23.2% |
| PKM2 P1 | 16.9% | 10.4% | 26.3% | 23.9% | 36.2% | 41.7% |
| PKM2 P2 | 20.0% | 13.3% | 26.0% | 24.6% | 48.5% | 48.0% |
| GDH G1 | 8.2% | 3.9% | 10.1% | 5.1% | 15.1% | 13.9% |
| GDH G2 | 6.1% | 1.4% | 9.7% | 5.3% | 12.5% | 9.8% |

Supplementary Table 10. Performance of the cryoPARES deep learning model (local refinement disabled) and the CESPED baseline model on the CESPED benchmark[3]. Best values in bold.

| | MAnE (°) | | wMAnE (°) | | MedAnE (°) | | $FSCR_{0.143}(V_0, V_1)$ (Å) | | $FSCR_{0.5}$(GT, V) (Å) | |
|---|---|---|---|---|---|---|---|---|---|---|
| EMPIAR ID | CESPED | cryoPARES | CESPED | cryoPARES | CESPED | cryoPARES | CESPED | cryoPARES | CESPED | cryoPARES |
| 10166 | 15.7 | **15.6** | 9.1 | **8.4** | 4.1 | **3.6** | 5.1 | **5.0** | 8.1 | **7.9** |
| 10786 | 32.6 | **30.1** | 29.5 | **27.1** | 7.3 | **7.1** | 3.8 | **3.5** | 7.6 | **7.0** |
| 10280 | **17.8** | 18.7 | **14.9** | 15.5 | 7.2 | **6.3** | 3.9 | 3.9 | 7.0 | **6.8** |
| 11120 | 44.7 | **42.6** | 41.1 | **39.1** | 7.0 | **5.2** | 4.1 | **3.9** | 8.3 | **8.0** |
| 10409 | 45.3 | **41.8** | 39.2 | **35.7** | 10.8 | **8.1** | 3.5 | 3.5 | 8.3 | **4.8** |
| 10374 | 35 | **34.1** | 24.8 | **24.2** | 5.8 | **5.2** | 3.7 | **3.5** | 6.5 | **5.9** |
| 10399 | 25.5 | **19.4** | 21.6 | **16.1** | 8.8 | **6.8** | 3.7 | **3.5** | 6.1 | **4.8** |
| 10648 | 13.3 | **12.9** | 10.6 | **10.2** | 12.2 | **10.1** | 3.8 | **3.7** | 6.5 | **5.7** |
| Simulated [illegible] | 6.0 | **4.6** | NA | NA | 5.7 | **4.9** | 4.5 | 4.5 | 4.8 | 4.8 |
| Consensus | 8.3 | **7.3** | 8.1 | **7.3** | 3.1 | **2.7** | 3.8 | **3.6** | 6.8 | **6.2** |
| Mean | 24.4 | **22.7** | 22.1 | **20.4** | 7.2 | **6.0** | 4.0 | **3.9** | 7.0 | **6.2** |
| std | 14.4 | **13.7** | 12.6 | **11.8** | 2.7 | **2.1** | 0.5 | 0.5 | **1.1** | 1.2 |

MAnE: mean angular error; wMAnE: weighted mean angular error; MedAnE: median angular error; $FSCR_{0.143}(V_0,V_1)$: reconstructed half-to-half FSC resolution at threshold 0.143; $FSCR_{0.5}$(GT,V): reconstructed to ground truth resolution at threshold 0.5. std: Standard deviation. Note that reconstructions for FSC estimation are performed using ground-truth shifts for comparison with the CESPED baseline model, that can only predict angular assignments.

Supplementary Table 11. Inference performance of cryo-FORUM, trained on the same apo datasets as cryoPARES

| Dataset | cryo-FORUM Median Geo Error (°) | cryoPARES (nnet only) Median Geo Error (°) |
|---|---|---|
| BGAL B1 | 9.8 | 4.0° |
| BGAL B2 | 14.2 | 6.0° |
| PKM2 P1 | 18.3 | 7.1° |
| PKM2 P2 | 17.4 | 7.0° |
| GDH G1 | 15.5 | 7.5° |
| GDH G2 | 16.2 | 7.2° |

Supplementary Table 12. Backbone RMSD between the apo state (training) and the liganded states (inference).

| | Apo vs Ligand 1 (Å) | Apo vs Ligand 2 (Å) |
|---|---|---|
| BGAL | 0.2 | 0.6 |
| PKM2 | 2.1 | 2.7 |
| GDH | 0.8 | 0.7 |
| TRPML1 | 0.4 | — |

Supplementary Table 13. Impact of dataset corruption in the GDH apo dataset on training quality, measured as a function of the G2 ligand inference quality. Two types of corruptions were tested: 1) mixing the experimental particles with white noise particles, and 2) randomizing the poses of a fraction of experimental particles.

| % of the dataset | FSC-half-to-half (0.143) Å | Median Geodesic error neural network (°) | Median Geodesic error after local refinement network (°) |
|---|---|---|---|
| Experimental vs $\mathcal{N}(0,1)$ | | | |
| 100% | 3.0 | 7.2 | 3.1 |
| 87.5% | 3.0 | 7.2 | 3.1 |
| 75% | 3.1 | 7.4 | 3.2 |
| 50% | 3.2 | 8.6 | 3.8 |
| Experimental vs Randomized angles | | | |
| 100% | 3.0 | 7.2 | 3.1 |
| 75% | 3.1 | 7.9 | 3.4 |
| 50% | 3.3 | 8.9 | 4.0 |

Supplementary Table 14 Training set size sensibility analysis performed on GDH. The inference statistics are computed using the GDH G2 dataset.

| Training set size | Validation loss | Validation mean geodesic error (°) | Validation median geodesic error (°) | Inference FSC (0.143, Å) | Nnet inference median geodesic error (°) | Nnet+local refinement inference median geodesic error (°) |
|---|---|---|---|---|---|---|
| 111K (100%) | 1.94 | 20.4 | 5.6 | 3.0 | 7.2 | 3.1 |
| 100K | 2.08 | 20.8 | 6.0 | 3.0 | 7.4 | 3.5 |
| 50K | 2.16 | 23.1 | 7.2 | 3.3 | 9.2 | 3.9 |
| 25K | 2.32 | 24.5 | 8.4 | 3.4 | 11.0 | 5.0 |
| 10K | 7.9 | 38.1 | 27.2 | NA | 21.2 | 15.4 |

Note: For the 10K example, the reconstruction was unsuccessful, hence, we do not report the FSC resolution here, as it will be misleading.

Supplementary Table 15. Performance comparison between the simple template matching local refinement algorithm (±6° step 2°), and the two-stage local refinement version (first ±6° step 2°, then ±2.1° step 0.7°).

| Dataset | Single-stage median Geodesic error | Two-stage median Geodesic error | Single-stage FSC | Two-stage FSC | RELION FSC |
|---|---|---|---|---|---|
| B1 | 1.6° | 1.0° | 2.8 Å | 2.6 Å | 2.4 Å |
| B2 | 2.2° | 2.1° | 3.2 Å | 2.9 Å | 2.6 Å |
| P1 | 3.1° | 2.9° | 3.5 Å | 3.5 Å | 3.4 Å |
| P2 | 2.9° | 2.2° | 3.3 Å | 2.9 Å | 2.8 Å |
| G1 | 3.4° | 2.2° | 2.9 Å | 2.7 Å | 2.7 Å |
| G2 | 3.1° | 2.0° | 3.0 Å | 2.9 Å | 2.8 Å |
| T1 | 3.7° | 2.8° | 2.7 Å | 2.6 Å | * 2.1 Å |

Note: * The T1 reconstruction was obtained using a multi-step cryoSPARC workflow rather than RELION.

### 3. Supplementary Notes

#### Supplementary Note 1: Data Augmentation

Each particle in a batch gets augmented with a probability of 0.95. Particles selected for augmentation are applied 1 to 4 rounds (uniformly random selection) of augmentation. In each round of augmentation, the following transformations are applied:

1. Random gaussian noise addition with a random standard deviation from 0 to 0.5 with probability 0.1.
2. Random uniform noise addition with a random scale from 0 to 2 with probability 0.2.
3. Random zoom-in of size 0% to 5% with probability 0.2.
4. Random gaussian blur addition with a random scale from 0 to 2.0 with probability 0.2.
5. Random erasing of patches of size 0% to 2% with probability 0.1.
6. Random elastic deformation with a scale of 10% and sigma of 10% with probability 0.1.
7. Random 90° rotation with probability 1.
8. Random rotation from −20° to 20° with probability 0.5.
9. Random shifts from −5% to 5% of the particle size with probability 0.5.

#### Supplementary Note 2: Model hyperparameters

Our architecture follows the image2sphere design[4] with the following implementation choices.

- Feature extractor: U-net with 5 encoder blocks and 4 decoder blocks. The number of channels in each encoder block is 16, 32, 64, 128 and 256. The kernel size is 5 pixels. Leaky ReLU is used as activation function. Upsampling is performed via bilinear interpolation. Batch normalization is used in each encoder and decoder block. The output is generated via a linear layer with 512 channels, leading to an output of shape Bx512xL/2xL/2.
- Image projector to S2: Default orthographic projector as proposed in the Image2Sphere paper with HEALPix grid order 3 (∼7.5°), where only 50% of the grid points are sampled. The feature map is projected from the 512 channels encoded in the Unet to 512 using a 1 x 1 Conv2d and then converted to spherical harmonics with $l_{max}$=12.
- $S^2$ convolution: 512 filters with global support on a HEALPix grid of order 4.
- SO(3) convolution: 64 filters with local support (angular range of $\pi/12$, 8 angles).
- Probability distribution discretization: HEALPix grid of order 4 (∼3.7°).

As encoder, in addition to the original ResNets and our selected U-net, other alternative architectures were tried:

- Group-equivariant U-net implemented with escnn, with up to 8 rotations.

- Neural operators in the form of U-shaped model[5].
- Conv-Mixer[6].

None of them beat the U-net in a consistent manner, with the Conv-Mixer being the second best and the Neural operator-based architecture the worse.

#### Supplementary Note 3: Particle pruning thresholds

There are several approaches to estimate direction-normalized robust z-score thresholds. For instance, they could be computed based on the expected fraction of bad picks given historical data. Here we estimate them by comparing the distribution of per-cone robust z-scores of the validation dataset against a dataset of "bad" particles (particles that were removed after 2D classification). Due to the significant number of false "good" particles in the validation set and false "bad" particles in the "bad" dataset, we model the distribution of neural network scores of both datasets as a Gaussian Mixture Model (GMM) with two components - one for "good" particles and one for "bad" particles. The threshold is then estimated as the intersection point between the density functions of the Gaussian distributions for "good" particles in the validation dataset and the density function for the bad particles in the "bad" particles dataset. The histograms and fitted GMMs used in this work can be seen at Supplementary Fig. 2.

#### Supplementary Note 4: Angular errors and local refinement

Supplementary Table 5 summarizes the results of a sensitivity analysis of the local refinement algorithm on sample PKM2 P2, varying the search range, angular step size, and filter cutoff. These results indicate that both having a narrower angular sampling and a higher-resolution low-pass filter help to push the resolution slightly further. However, these gains saturate rapidly. This evidence, alongside the FSC curves in the Main Text showing that search ranges beyond 4° do not yield significant gains, suggests that the network's initial pose accuracy—limited in part by the hyperparameter that defines the resolution of the network's SO(3) grid—is an important factor defining the optimal refinement window, rather than the filter cutoff alone.

#### Supplementary Note 5: Running time

Supplementary Table 6 shows the processing throughput, in particles $min^{-1}$ $GPU^{-1}$, for cryoPARES neural network inference and the local refinement step with different search range values using two different GPUs. As can be seen in Supplementary Table 7, cryoPARES throughput is comparable to cryoSPARC homogeneous-refinement and at least one order of magnitude faster than RELION. To explore the effect of symmetry in running times, two additional datasets from EMPIAR[7] were also studied: EMPIAR-10280 and EMPIAR-10166.

We also explored the computational efficiency of performing a grid-based SO(3) search using a high-resolution apo-structure. However, our results indicate that this remains significantly more computationally expensive than the amortized inference provided by cryoPARES.

Quantitatively, a global alignment in RELION at HEALPix level 4 (1.8° sampling with oversampling) requires evaluating approximately 121 million poses per particle. In our tests using a PKM2 dataset of 125K particles on 4x A5000 GPUs, this search took 2 hours and 38 minutes for a single iteration. Even reducing the sampling density to HEALPix level 3 (3.7° sampling, ~23M poses with oversampling) required 1 hour and 24 minutes. In contrast, cryoPARES pose estimation (neural net + local refinement with ±4° step 2°) on the same hardware took 7 minutes.

The difference is even more evident when considering computational complexity. We estimated that a global search at HEALPix level 4 costs ~20 TFLOPs per particle, or ~3 TFLOPs when using HEALPix level 3, whereas the cryoPARES inference pass plus local refinement costs only ~0.06 TFLOPs. Furthermore, we observed that an auto-refinement job starting from a 6 Å low-pass filtered map took 59 minutes, close to the 67 minutes required using a 30 Å low-pass filtered map. This behaviour may reflect an inherent trade-off in traditional hierarchical approaches: achieving high-resolution alignment would require a very fine angular search in the initial global step, which is computationally expensive, whereas using a coarser angular sampling (e.g., ~15°) limits the resolution of the first reconstructed map. As a result, even when starting from a higher-resolution reference, the first iteration may converge to a lower-resolution structure that is then used in subsequent refinement steps, reducing the practical benefit of the higher-resolution initialization.

#### Supplementary Note 6: Automatic particle pruning results

The cryoPARES pruning score quantifies angular assignment confidence rather than image quality in an absolute sense. As a consequence, the set of rejected particles is expected to be heterogeneous: it will include genuine junk particles (aggregates, ice contamination, carbon edge picks) as well as morphologically intact protein particles for which the network could not produce a confident pose estimate, for example due to low signal-to-noise ratio or projection directions underrepresented in the training data.

To characterize the rejected particles directly, we performed a 2D classification job in RELION on the particles discarded by cryoPARES in the TRPML1 T1 dataset. Inspection of the resulting class averages confirms this expectation: while the majority of classes show featureless or heterogeneous density consistent with junk particles, at least one class genuine TRPML1 particles (see Supplementary Fig. 5).

To verify that this rejection is nonetheless well-founded from an angular accuracy standpoint, we compared the angular errors of the cryoPARES-retained particles against those of the rejected particles that fall into this good 2D class—i.e., the subset that is morphologically recognizable as genuine TRPML1 but was nevertheless pruned. Using the RELION auto-refine poses as a reference, we measure a median angular error of 3.7° for the retained particles, compared to 26.7° for the rejected particles in the good 2D class. This approximately seven-fold difference confirms that the rejected genuine particles carry substantially less reliable orientation information than those retained. Their inclusion would therefore introduce angular noise into the reconstruction, and their removal is well-justified by the direction-normalized confidence score described in Methods Section 4.10 and Supplementary Note 3. This demonstrates that cryoPARES does not exclusively reject junk; it also rejects particles whose pose estimates fall below the confidence threshold, regardless of their structural integrity.

#### Supplementary Note 7: Dealing with model bias

The core idea behind cryoPARES is that the structural features that guide particle alignment in an apoprotein are largely preserved when small molecules bind to it. This hypothesis is based on the observation that ligand binding often induces localized conformational changes while maintaining the protein's overall shape, which is the crucial part for alignment. However, this assumption may not hold in cases where ligand binding triggers substantial conformational changes. In such scenarios, two potential sources of bias need careful consideration: model bias from the neural network predictions could lead to incorrect density interpretations, and template bias could arise when using high-resolution apo structures as references during local refinement[2].

For ligand determination, template bias presents fewer concerns because ligands are absent from the reference apo structure[2]. False positives, such as endogenous lipids occupying binding sites and being misinterpreted as ligand densities, can generally be resolved by improving map resolution or by excluding these components from the reference model during local refinement. However, the false negative case, in which a ligand-bound structure is not resolved either due to model bias or insufficient reconstruction quality, could occur and should be detected[2,8–10], to ensure that structural screenings are reliable.

Our approach to detect model bias is based on applying the gold standard for the local refinement step while using a medium-resolution reference (by default, low-pass filtered at 6 Å). If the half-to-half FSC resolution does not improve substantially beyond the filtering resolution, we consider the density to be affected by model bias or to be of low quality and cryoPARES recommends users to run a classical reconstruction program to rule out a false negative case.

Supplementary Fig. 6 illustrates the detection of two extreme cases. In the first case, we calculate a local refinement (4°) on a dataset with randomised orientations, simulating the complete failure of the neural network to infer poses (Supplementary Fig. 6 a,b). In the second case, we simulate the effect of high model bias by creating an Einstein from noise map that matches the training protein and running local refinement after (Supplementary Fig. 6 c,d). As shown in Supplementary Fig. 6 g, using a mid-resolution, 6Å template for local refinement (instead of a high-resolution template) easily identifies reference bias in both cases by the FSC curves quickly dropping to zero at the template's resolution. Conversely, if a high-resolution template is used instead, we obtain well-defined high-resolution structures with high FSC resolution that would lead to incorrect structural interpretation. When orientations correctly predicted by cryoPARES are used instead, the usage of a reference low-pass filtered at 6 Å does not prevent the reconstruction from reaching a resolution significantly better than the filtering cut-off (Supplementary Fig. 6 e,f).

In order to quantify the amount of model bias in our reconstructions, we also computed the Ω metric proposed by Lucas et al.[2]:

$$\Omega = \frac{\sum_{i \in \mathbf{M}} \mathbf{V}_i^{\mathrm{full}} - \sum_{i \in \mathbf{M}} \mathbf{V}_i^{\mathrm{omit}}}{\sum_{i \in \mathbf{M}} \mathbf{V}_i^{\mathrm{full}}} \quad (1)$$

where

$$\mathbf{M} = \{i : (\mathbf{T}_i^{\mathrm{full}} - \mathbf{T}_i^{\mathrm{omit}}) > \mathrm{threshold}\} \quad (2)$$

Here $\mathbf{V}^{\mathrm{full}}$ is the volume reconstructed after aligning with the full template ($\mathbf{T}^{\mathrm{full}}$), $\mathbf{V}^{\mathrm{omit}}$ is the volume reconstructed after aligning with the template with omitted density ($\mathbf{T}^{\mathrm{omit}}$) and $\mathrm{threshold}$ is a value defined by default as 10% of the max value of ($\mathbf{T}_i^{\mathrm{full}} - \mathbf{T}_i^{\mathrm{omit}}$). As in the original publication, we employed full templates simulated from the atomic models, and omit templates were generated by random removal of 3% of the residues.

In this work, we also compute a slightly modified version to account for cases where regions in $\mathbf{V}^{\mathrm{omit}}$ show higher density than $\mathbf{V}^{\mathrm{full}}$ (negative differences), which can mathematically cancel out regions of true template bias (positive differences). This cancellation effect can lead to underestimation of actual template bias, as it combines two distinct phenomena in the calculation. While positive differences, where the full reconstruction shows higher density than the omit reconstruction, likely represent genuine template bias where the full template has influenced the reconstruction, negative differences could arise from various sources unrelated to template bias, such as local reconstruction variations, noise in the reconstruction process, differences in particle alignment, or natural flexibility in the structure. To address this limitation, we propose to set all negative differences to zero, effectively focusing only on regions where the full reconstruction shows higher density:

$$\Omega^0 = \frac{\sum_{i \in \mathbf{M}} \max(\mathbf{V}_i^{\mathrm{full}} - \mathbf{V}_i^{\mathrm{omit}}, 0)}{\sum_{i \in \mathbf{M}} \mathbf{V}_i^{\mathrm{full}}} \quad (3)$$

Supplementary Table 9 shows that the amount of quantified template-bias for cryoPARES is much smaller for datasets which had poses inferred by cryoPARES prior to local refinement compared to datasets which had been assigned random orientations. It is also clear that, for most cases, the use of a template which has not been low-pass filtered can increase the amount of template bias in the reconstruction, and it seems that this effect is more pronounced for the target with worse resolution, PKM2. This is partially caused by the fact that the value of $\Omega$ is highly correlated with the resolution of the map. We have measured that the Pearson's correlation coefficient between the cryoPARES Unfiltered $\Omega$ column of Supplementary Table 9 and the $\mathbf{V}^{\text{full}}$ resolution is of 0.9.

In addition to the model bias detection strategy, cryoPARES implements several mechanisms for model bias prevention. The model is trained on independent half-sets of the training particles using extensive data augmentation, resulting in two independently trained networks that are used at inference time. While this does not constitute a proper gold standard since the models are used to infer the poses of other samples, the usage of two uncorrelated models minimizes the chances of systematic biases in half maps, preventing FSC resolution inflation. Furthermore, the architecture includes a dropout-like layer that remains active during both training and inference to introduce variability in the predictions and reduce the risk of overconfident, deterministic outputs driven by memorized structural features. The local refinement template is low-pass filtered at 6 Å by default, which prevents model bias at the resolutions needed for high-resolution on-the-fly ligand screening experiments. CryoPARES further differs from traditional refinement approaches in that it does not estimate particle shifts during initial stages, but only angular assignments. High-resolution features can only emerge during the local refinement stage, where orientations and shifts are jointly optimized. Consequently, any high-resolution feature arises from this refinement step and can therefore be validated using gold-standard FSC. Particle weights used at reconstruction are based on both the neural network confidence and the significance of the cross-correlation score compared to neighbouring orientations, meaning that when the best orientation is not significantly better than others, the particle's contribution is smaller. Finally, an optional step of particle pruning via direction-normalized scores based on cross-correlation has been implemented, which can be used in a similar manner to the SNR threshold used by Lucas et al.[2].

#### Supplementary Note 8: Comparison with other approaches

We evaluated the performance of the cryoPARES deep learning model using the CESPED benchmark[3], comparing it directly against the original CESPED baseline model. The CESPED benchmark consists of 10 diverse cryo-EM targets, where each dataset is divided into two independent half-sets. Following the benchmark protocol, the models are trained on one half-set, and inference is subsequently performed on the remaining subset. The predicted angular assignments are then compared against the ground-truth (GT) labels provided by the benchmark. The primary goal of this comparison is to assess the precision of the neural network's angular assignments and determine how these improvements translate into the quality of the resulting 3D reconstructions.

As shown in Supplementary Table 10, the cryoPARES deep learning model consistently outperforms the baseline across all evaluated metrics. While the mean angular error (MAnE) provides a general overview, it is often skewed by a small fraction of highly misaligned particles that act as outliers. To provide a more robust measure of the model's central performance, we have included the median angular error (MedAnE). Across the benchmark, cryoPARES achieves an average MedAnE of 6.0°, a 1.2° improvement over the baseline's 7.2°. In specific cases, such as EMPIAR 10409 and 11120, the median error improves by approximately 25%, demonstrating a significant boost in precision for the majority of the particle population.

Beyond angular accuracy, the cryoPARES deep learning model shows consistent improvements in reconstruction resolution as measured by the Fourier Shell Correlation (FSC). The resulting improvements in the $FSC_{0.5}$(GT, V) metric, reaching up to 3.5 Å in specific targets, underscore that the modifications in the cryoPARES deep learning model led to relevant gains in structural detail.

It is also important to highlight a fundamental functional difference: while the CESPED baseline is a machine learning prototype limited strictly to angular estimation, cryoPARES is a comprehensive pipeline capable of shift estimation, local refinement, and full volume reconstruction. However, for the purposes of this test, only the global angular estimation capability of cryoPARES is being employed.

To situate the performance of cryoPARES within the broader landscape of amortized pose inference, we compared our method against cryo-forum[11] and cryoFIRE[12]. These methods, like cryoPARES, aim to predict particle orientations directly from images using neural networks.

Across all evaluated datasets, cryo-FORUM yields median geodesic errors ranging from 9.8° to 18.3° (Supplementary Table 11). In contrast, cryoPARES consistently achieves substantially lower errors, typically in the <7° range without local refinement. This corresponds to an approximate two-fold reduction in median error.

We also sought to include cryoFIRE[12] in this benchmark. However, we found the training process to be less robust for these specific datasets; the model failed to converge across several hyperparameter configurations. This highlight's one of the key advantages of the cryoPARES architecture: its increased stability and ease of use when applied to diverse experimental datasets.

##### Supplementary Note 9: Tolerance to conformational changes

To evaluate the sensitivity of cryoPARES to conformational variability, we computed backbone RMSDs between the apo structures used for training and the ligand-bound structures used for inference (Supplementary Table 12). Across all targets, these deviations range from minimal (e.g., BGAL: 0.2–0.6 Å; TRPML1: 0.4 Å; GDH: 0.7–0.8 Å) to moderate (e.g., PKM2: 2.1–2.7 Å).

Overall, cryoPARES remains robust in the presence of moderate conformational changes (see Supplementary Table 12) . In particular, PKM2 shows backbone deviations of up to 2.7 Å while still yielding successful reconstructions. Notably, the P1 state, despite having a smaller RMSD than P2, results in lower resolution, indicating that reconstruction quality does not correlate directly with the magnitude of structural deviation.

##### Supplementary Note 10: Requirements of the training set

In order to train cryoPARES, it is necessary to employ a training set of sufficient quality. Based on the experiments included in this work, we advise using for training at least 100K pre-aligned particles capable of yielding a high-resolution map (at least, 3.5 Å). We also recommend studying the angular distribution of the pre-aligned particles and the anisotropy of the map (for example, measuring the FSO[13], as the biases in the training set can be translated to biases in the inferred poses.

As it can be seen in Supplementary Table 13, cryoPARES is quite robust to the presence of corrupted particles, or even incorrectly aligned particles. However, when the fraction of such errors becomes a large portion of the training set, it can have a severe impact on the performance. In order to reduce this impact, we suggest employing angular consensus[14] from two map refinements (e.g. two RELION auto-refinement runs), discarding those particles in which the alignment deviates by more than 5° between the two refinements.

With respect to training size requirements, Supplementary Table 14 shows that accuracy degrades slowly and smoothly as the training set size decreases, with a noticeable drop under <50K particles. In practice, most of our experiments use 100K–200K particles per model, which provides a good balance between accuracy and computational cost.
